# H3K27me3 and H2A.Z prime cold regulated genes, and their remodelling governs plant cold response

**DOI:** 10.64898/2026.04.16.718857

**Authors:** Sarah Mermet, Sarah Muniz Nardeli, Simon Amiard, Aline V Probst, Peter Kindgren

## Abstract

To elucidate the contribution of chromatin modifications in plant cold response, we performed ChIP-seq for H3K27me3 and H2A.Z in Arabidopsis exposed to short-term cold. We combined our epigenetic data with NET-seq to investigate the direct transcriptional effects of histone marks. Prior to any stress cue, cold regulated genes share a similar chromatin environment with high H2A.Z and H3K27me3. H3K27me3 levels do not correlate with transcriptional activity or elongation speed. However, REF6-mediated reduction of H3K27me3 is essential for regulation of cold controlled genes. H2A.Z occupancy changes revealed a negative correlation between cold-induced changes to H2A.Z and RNAPII activity at differentially expressed genes. Importantly, changing H2A.Z levels preceded transcriptional changes, indicating that the variant functions as a critical cold-induced switch. Further, our data suggests that high H2A.Z levels slow down RNAPII. Thus, H2A.Z is essential for the transcriptional response and a decreased H3K27me3 level is important for the genomic adaptation to cold.

## INTRODUCTION

DNA in the nucleus is wrapped around histone octamers, which are composed of histones H2A, H2B, H3 and H4 to form the nucleosomes, the basic subunits of chromatin^1^. Heterochromatin, or condensed chromatin, contains transcriptionally repressed genomic regions, while euchromatin rather contains transcriptionally permissive regions^2,3^. Chromatin compaction and regulation of transcription are controlled by epigenetic marks, such as post-translational modifications of histones or incorporation of non-canonical histone variants^4,5^. For example, acetylation of histone H3 decreases its positive charge, thereby reducing its interaction with negatively charged DNA and inducing more open chromatin^6,7^. Other modifications associated with active transcription include H3K4me2, H3K4me3, H3K36me3 or H2Bub. In contrast, H3K27me3 set by Polycomb Repressive complex 2 (PRC2) and H3K9me2 are associated with more compact chromatin and are involved in transcriptional repression^8,9^. Thus, histone tail modifications are generally classified as either transcriptionally activating or repressive^8^. However, most of our understanding about the chromatin environment and its effect on transcriptional activity comes from studies in cell cultures and plants grown under constant conditions. Temperature fluctuations, such as cold, change the dynamics of biochemical processes (e.g. transcription), but we know relatively little about how the chromatin signature adjusts to novel conditions and stress cues.

The enrichment of certain marks differs between mammals and plants, such as H3K27me3^10^. In mammals, H3K27me3 is enriched at the promoters of non-transcribed genes associated with facultative heterochromatin. In contrast, in plants, H3K27me3 is present throughout the gene body, with higher abundance towards the Transcription Start Site (TSS)^10^. In Arabidopsis, it has been shown that a subset of cold response genes, transcriptionally reprogrammed at low temperature to promote cold acclimation and stress adaptation, loose H3K27me3 during the days following exposure to cold, although this loss was assessed only at multi-day time points and therefore does not reflect the rapid transcriptional induction observed within hours; notably, the reduced H3K27me3 levels persist after plants are returned to 22°C, even though these genes are rapidly repressed again, indicating that H3K27me3 is not essential for their silencing^11^. In addition, H3K27me3 undergoes redistribution after 3 h of cold exposure, although the correlation with transcription at this stage is weak, and the next time point examined was 3 days shows a similar pattern, leaving the intermediate early phase uncharacterized^12^. These studies point to a role for H3K27me3 in the cold response in plants, but the extent of H3K27me3 changes in response to short-term cold exposure and their contribution to the transcriptional cold response remain largely unknown.

Among histone variants, H2A.Z is highly conserved in all eukaryotes and has been extensively studied^13–16^. Incorporation of H2A.Z alters chromatin accessibility; it increases nucleosome stability^17^ with a stronger binding to DNA^18^, but also significantly increases H2A.Z-H2B dimer dissociation^17^. Its involvement in transcription regulation has been demonstrated, but its role varies depending on the organism and/or context studied^19–21^. Generally, H2A.Z is highly enriched at the +1 nucleosome, the first nucleosome after the TSS, and thereby involved in the transcriptional barrier generated by the +1 nucleosome^22–25^. It has also been shown that H2A.Z is enriched at the gene body of transcriptionally inactive genes^26–28^. In addition to its genomic distribution, the regulatory function of H2A.Z is further modulated by post-translational modifications, adding an additional layer of chromatin regulation. H2A.Z acetylation is generally associated with transcriptional activation^29^, whereas mono-ubiquitination correlates with transcriptional repression^27^, underscoring the context-dependent roles of H2A.Z at regulatory nucleosomes. In Arabidopsis, H2A.Z has been shown to play a role in the perception of ambient temperature^18,30,31^. Interestingly, nucleosomes of temperature-responsive genes are enriched in H2A.Z at non-inducible temperatures, which likely confers inducible gene expression and higher response dynamics^30^. Thus, the essential role of H2A.Z in temperature adaptation of eukaryotic cells is well established, but because of its enrichment at both active and in the gene body of inactive genes, many open questions remain.

To gain a better understanding of changes in the chromatin environment, we performed ChIP-seq for H3K27me3 and H2A.Z in response to short-term exposure to cold (3 and 12 hours at 4°C). We identified a common chromatin signature (high H3K27me3 and H2A.Z levels) for both cold up- and down-regulated genes prior to cold stress, indicating a possible key stress responsive chromatin environment in Arabidopsis, already at control conditions. Further, by combining the epigenomic approach with Native Elongation Transcript sequencing (NET-seq)^32^, we identified a negative correlation between changing H2A.Z levels and RNA Polymerase II (RNAPII) activity in response to cold, with corresponding changes to the kinetics of RNAPII elongation over nucleosomes. We find that H2A.Z occupancy reduction precedes RNAPII activity at cold-sensitive genes, indicating the variant as an essential transcriptional cold-induced switch. In contrast, changes in H3K27me3 levels did not correlate with transcriptional activity nor kinetics. Nevertheless, we show that H3K27me3-demethylation activity by RELATIVE OF EARLY FLOWERING 6 (REF6) is necessary for proper up- and down-regulation in response to cold of its cold-regulated target genes.

Thus, our study shows that changing levels of the repressive histone mark H3K27me3, induced by cold, is independent of transcription activity and that cold induced changes in nucleosome composition by eviction of the histone variant H2A.Z, in contrast, is a prerequisite for changes in transcriptional activity. Overall, our results suggest a necessary re-definition of what a repressive or activating chromatin signature entails.

## RESULTS

### Transcription activity correlates with changes in H2A.Z but not H3K27me3 in response to cold

To understand how the chromatin landscape changes in response to short term cold exposure, we performed ChIP-seq for a histone mark (H3K27me3), and a histone 2A variant (H2A.Z) in 10-day-old seedlings grown in long day conditions at 22°C and exposed to 4°C for 3 and 12h. Prior to cold treatment, the levels of the mark and histone variant along the gene body of expressed and not expressed transcription units showed the expected pattern, corroborating that our data was robust. H2A.Z levels were enriched at the +1 nucleosome for expressed genes (FPKM>1 at 22°C in plaNET-seq^32,33^) and throughout the gene body for non-expressed genes (FPKM<1; Figure 1a). The H3K27me3 levels were spread-out along the gene body and enriched in non-expressed genes (Figure 1b). Exposure to cold (3 or 12 hours at 4 °C), revealed few changes for either expressed or non-expressed genes regarding H2A.Z (Figure 1a) or H3K27me3 (Figure 1b). Furthermore, when focusing on genes with significant changes in H3K27me3 or H2A.Z levels during cold treatment, a poor overlap was observed, whether considering gene-level differential enrichment (fold change>2) or peak-level changes in signal intensity (Δintensity>1.5xSD) (Supplementary Figure 1a-d, Supplementary Tables 1 and 2).

**Figure 1.**
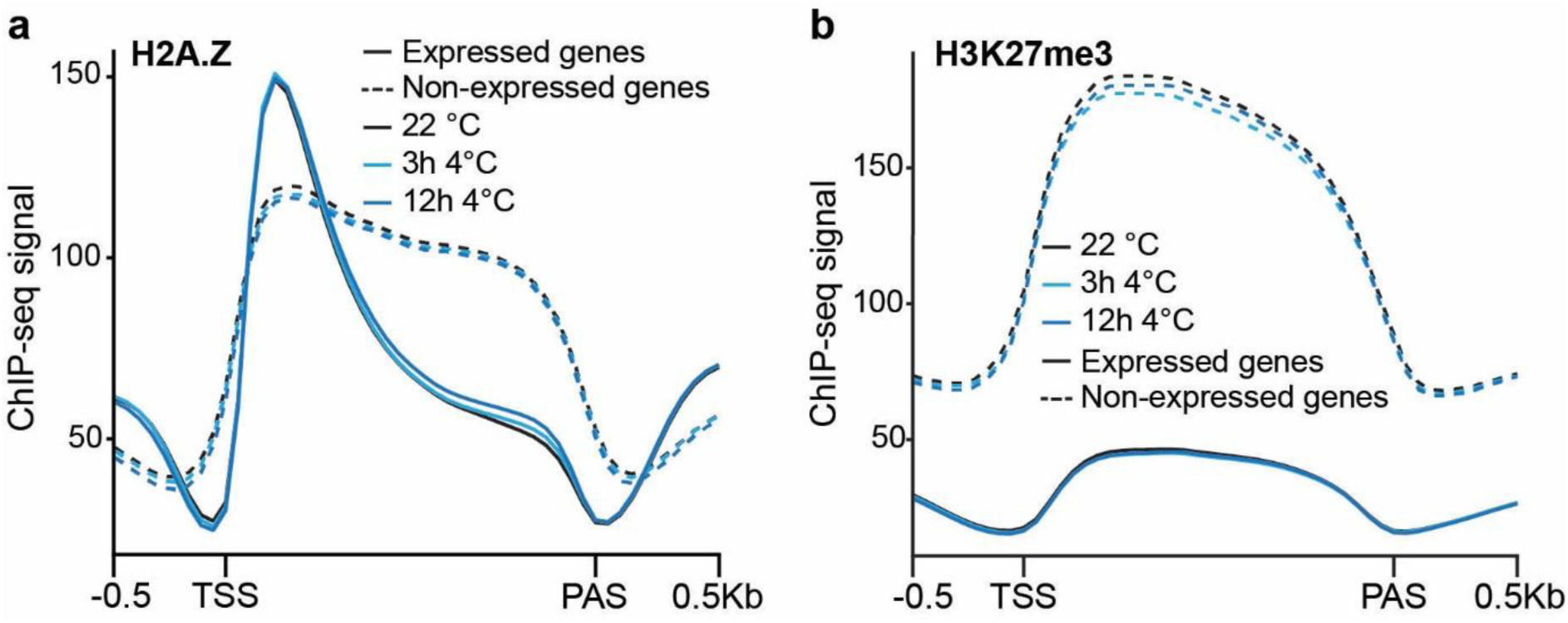
Levels of H2A.Z and H3K27me3 along gene bodies in Arabidopsis a-b) Metagene plots of ChIP-seq mean signal of H2A.Z **(a)** or H3K27me3 (**b**). Expressed genes are represented by solid lines and non-expressed genes by dashed lines; colours indicate temperature: 22°C in black, 3h at 4°C in light blue and 12h at 4°C in dark blue. The metagene plot covers the gene body and includes 500 bp flanks upstream and downstream of TSS and PAS, respectively. TSS: Transcription Start Site, PAS: Poly(A)-Signal.

To establish a link between the chromatin state changes of H3K27me3 and H2A.Z and differential expression induced by cold, we used previously published plaNET-seq data in similarly grown and cold treated seedlings as our ChIP-seq experiment. The advantage of using plaNET-seq data compared to a steady state level RNA technique (i.e. RNA-seq) is that plaNET-seq directly measures transcriptional activity on the DNA template, providing a more direct view of transcription dynamics in relation to histone modifications. First, we used differentially expressed genes (DEGs) based on plaNET-seq^32^ at 3h and 12h 4°C compared to 22°C. At 3 h, 3222 genes were down-regulated and 3697 genes up-regulated, whereas at 12 h, 3882 and 3936 genes were down- and up-regulated, respectively. Next, we examined how changing levels (intensity) of gene body H3K27me3 and H2A.Z (Supplementary Table 3) correlated overall with transcriptional activity in response to cold temperatures. We detected a negative correlation between transcriptional activity and H2A.Z levels in differentially expressed genes (Figure 2a-b). A substantial overlap was shown between differentially expressed genes and genes showing changes in gene body H2A.Z levels at both 3h and 12h at 4°C, with H2A.Z loss predominantly associated with gene activation (383/696 genes at 3h; 993/1493 at 12h) and H2A.Z gain with repression (104/237 at 3h; 862/1817 at 12h, Supplementary Figure 2a-d, Supplementary Tables 4-5). Examples of the change in gene body H2A.Z level and the correlated transcriptional response can be seen in Figure 2c-d. In contrast, we could not detect any correlation between changing transcription activity levels and H3K27me3 levels (Figure 2e-f), as illustrated by Venn diagrams comparing DEGs with genes showing changes in histone mark levels (Supplementary Figure 2e-h, Supplementary Tables 4-5). However, we detected more genes with reduced H3K27me3 levels (711 genes after 3h 4°C and 447 genes after 12h 4°C) compared to enriched (14 genes after 3h 4°C and 173 genes after 12h 4°C). Examples of genes that are up- or down-regulated but depleted in gene body H3K27me3 can be seen in Figure 2g-h. Taken together, our data indicates that the H3K27me3 mark does not correlate with transcriptional changes induced by cold treatment. In contrast, changing gene body H2A.Z levels at cold-responsive DEGs negatively correlates with their transcriptional activity.

**Figure 2.**
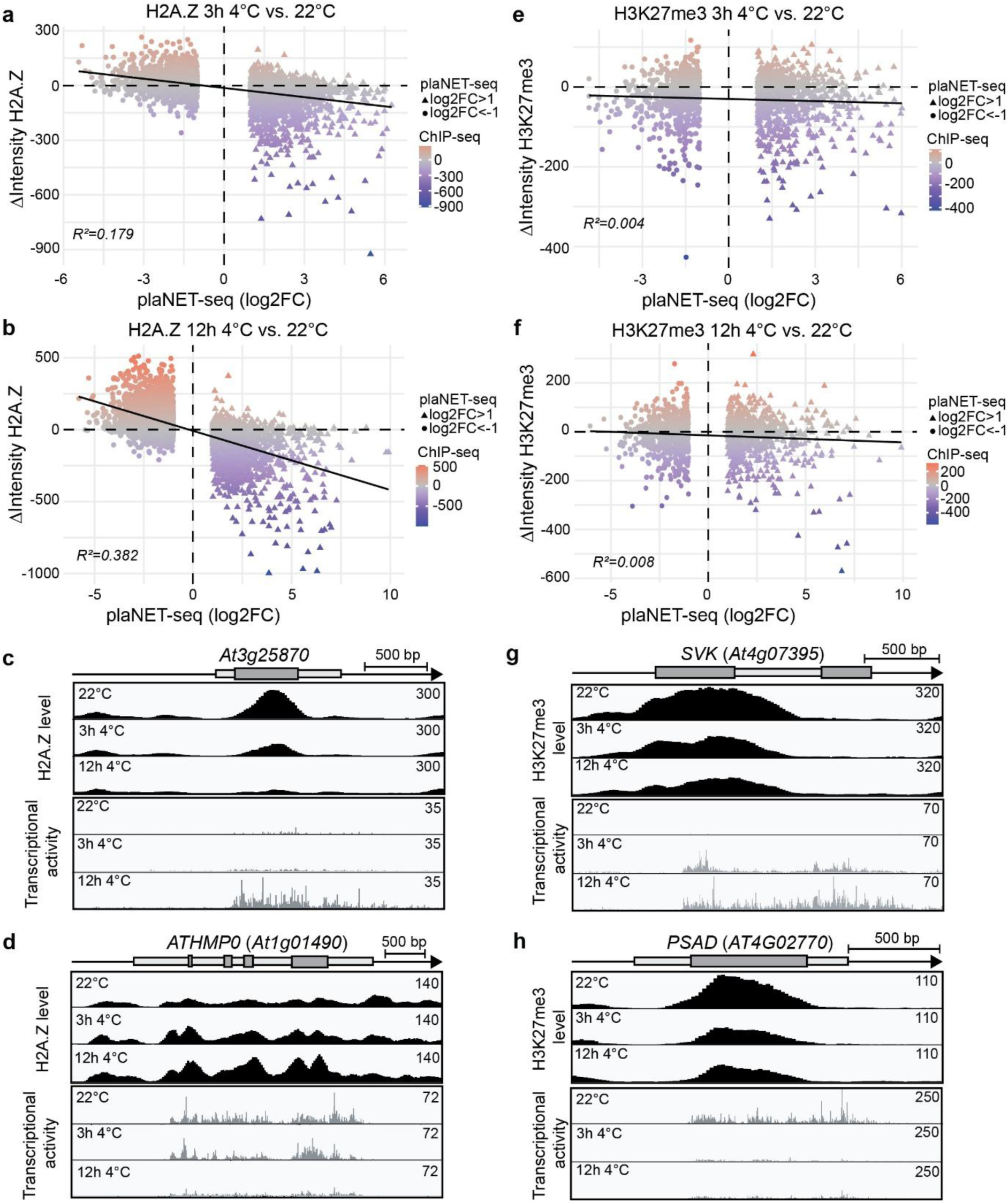
Correlation of H2A.Z and H3K27me3 levels to transcriptional activity after cold exposure. a-b, e-f) Correlation plots between differentially transcribed genes in plaNET-seq (log₂ fold change; upregulated (log_2_FC>1): triangles, downregulated (log_2_FC<-1): circles) and changes in ChIP-seq signal intensity for gene body H2A.Z **(a–b)** or gene body H3K27me3 **(e–f)** after exposure to 4 °C for 3 h **(a, e)** or 12 h **(b, f)**, relative to 22 °C. Each point represents an individual ChIP-seq peak. Correlations were calculated using a linear regression model; solid lines indicate the fitted regression, and the coefficient of determination (R²) is shown on each plot. Colours indicate the magnitude of the ChIP-seq signal difference (Δ intensity). **c-d, g-h)** Examples of cold-responsive genes, up-regulated in cold: At3g25870 **(c)** and SVK, At4g07395 **(g)** or down regulated in cold: ATHMPO, At1g01490 **(d)** and PSAD-1, At4g02770 **(h)**. Screenshots are from ChIP-seq H2A.Z **(c-d)** or H3K27me3 **(g-h)** and plaNET-seq datasets. Higher occupancy of H2A.Z or H3K27me3 and elevated transcriptional activity are indicated by higher peak density and amplitude. At3g25870 and ATHMPO loose or gain H2A.Z, respectively, upon cold exposure, while SVK and PSAD-1 loose H3K27me3.

### Cold responsive genes have a distinct chromatin signature at 22°C

We then delved deeper into the cold regulated genes to understand whether their chromatin environment prior to the cold exposure could pre-determine their transcriptional response. To avoid rapid and transient effects of the cold response, we focused on genes differentially regulated after 12h at 4°C. When plotted, we observed that downregulated genes after 12h at 4°C had high transcriptional activity prior to the cold treatment and upregulated genes low transcriptional activity (Figure 3a).

**Figure 3.**
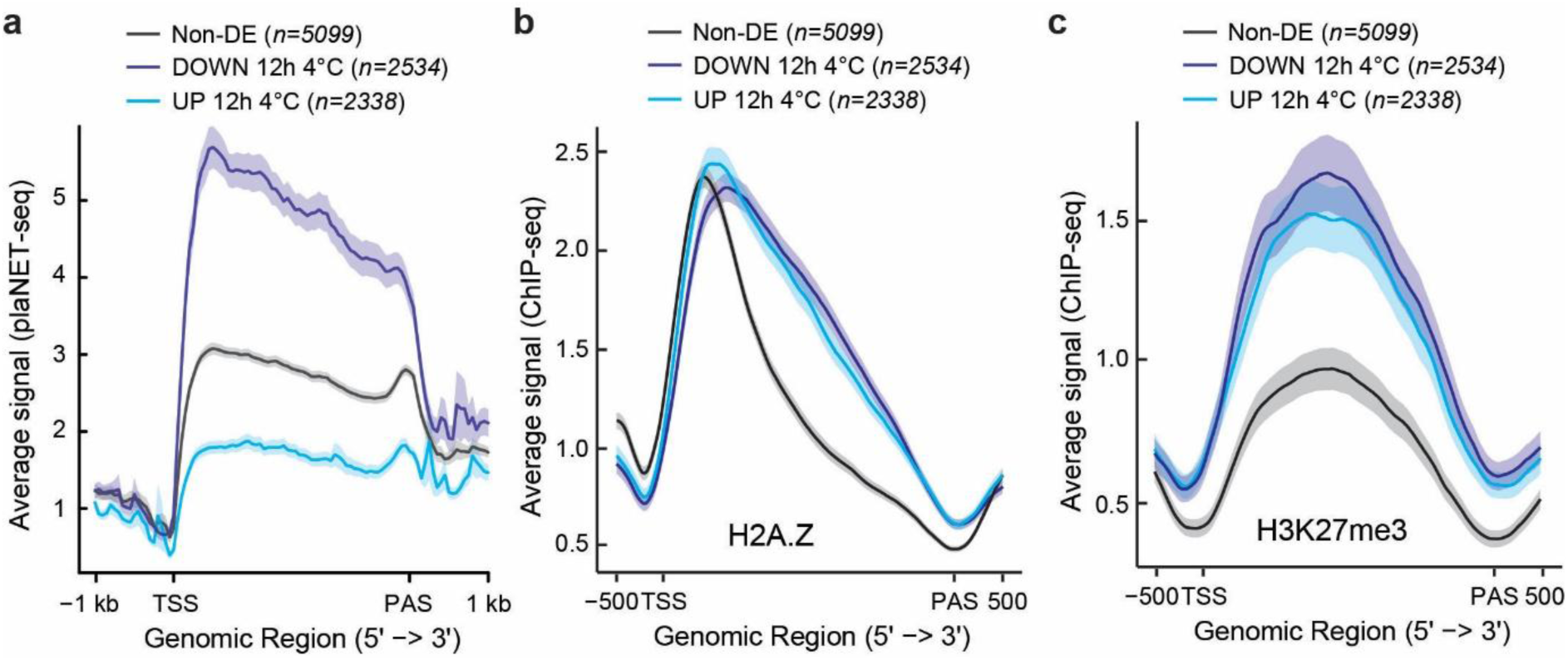
Cold-regulated genes have a distinct chromatin signature at 22°C. a-c) Metagene plots of average signal from plaNET-seq **(a)** or ChIP-seq of H2A.Z **(b)** or H3K27me3 **(c)** at 22°C of expressed non-regulated genes in cold (non-DE, in black, n=5099), down regulated (DOWN 12h 4°C, in dark blue, n=2534) or up regulated (UP 12h 4°C, in light blue, n=2338) genes in plaNET-seq at 12h at 4°C. The metagene plot covers the gene body and includes 500 bp flanks upstream and downstream of TSS and PAS, respectively. TSS: Transcription Start Site, PAS: Poly(A)-Signal. The shaded area shows a 95% confidence interval for the mean.

Surprisingly, when we plotted H3K27me3 and H2A.Z levels, genes differentially regulated during cold showed higher levels of both gene body H3K27me3 and H2A.Z at 22°C, which corresponded to poor correlation between mark/variant level and transcriptional activity (Figure 3b-c, Supplementary Figure 3a-b). We further corroborated these results with other available ChIP-seq data sets^12,34,35^ (Supplementary Figure 3c-e). Our data clearly shows that cold-responsive genes have a distinct chromatin environment compared to genes that do not respond to cold temperatures, prior to the cold exposure. Thus, our data is questioning a simplistic repressive/activating role for H3K27me3 and H2A.Z in influencing transcriptional rates, rather, we could identify a cold-responsive chromatin signature already at 22°C.

### Changing H2A.Z levels precede transcription changes and are important for proper gene regulation in response to low temperature

At differentially expressed genes, transcriptional changes in response to cold correlate with changes in the composition of the +1 and internal nucleosomes (i.e. nucleosomes within the gene body, excluding the +1 nucleosome). At the +1 nucleosomes, H2A.Z showed gain and loss at down- and up-regulated genes, respectively (Figure 4a-c). A similar trend was evident at internal nucleosomes (Figure 4d-f), indicating that the levels of H2A.Z change along the whole gene body. The general negative correlation between H2A.Z levels and transcriptional activity (Figure 2) led us to consider two possible hypotheses. Either cold-induced changes in H2A.Z levels are a consequence of altered transcriptional activity or changes in H2A.Z levels are a prerequisite for gene induction or repression. To test these hypotheses, we selected specifically differentially expressed (DE) genes after 12h at 4°C (i.e. genes that were DE at 12h 4°C with a FC>2 but not DE at 3h 4°C with a FC<1.5, Supplementary Table 6) and analysed H2A.Z levels between 22°C and after cold exposure. The plaNET-seq expression profiles of the selected genes at 22°C, 3h 4°C and 12h 4°C are shown in Supplementary Figure 4. For upregulated genes, a significant drop in H2A.Z levels was detected already at 3h 4°C, prior to a change in transcriptional activity (Figure 5a-c), as exemplified by *RESPONSIVE TO DEHYDRATION 19* (Figure 5c). A similar pattern was detected for downregulated genes, the increase of H2A.Z levels preceded the decrease in transcriptional activity at 12h 4°C (Figure 5d-f), albeit a small downregulation trend could be seen at 3h 4°C (Supplementary Figure 4b).

**Figure 4.**
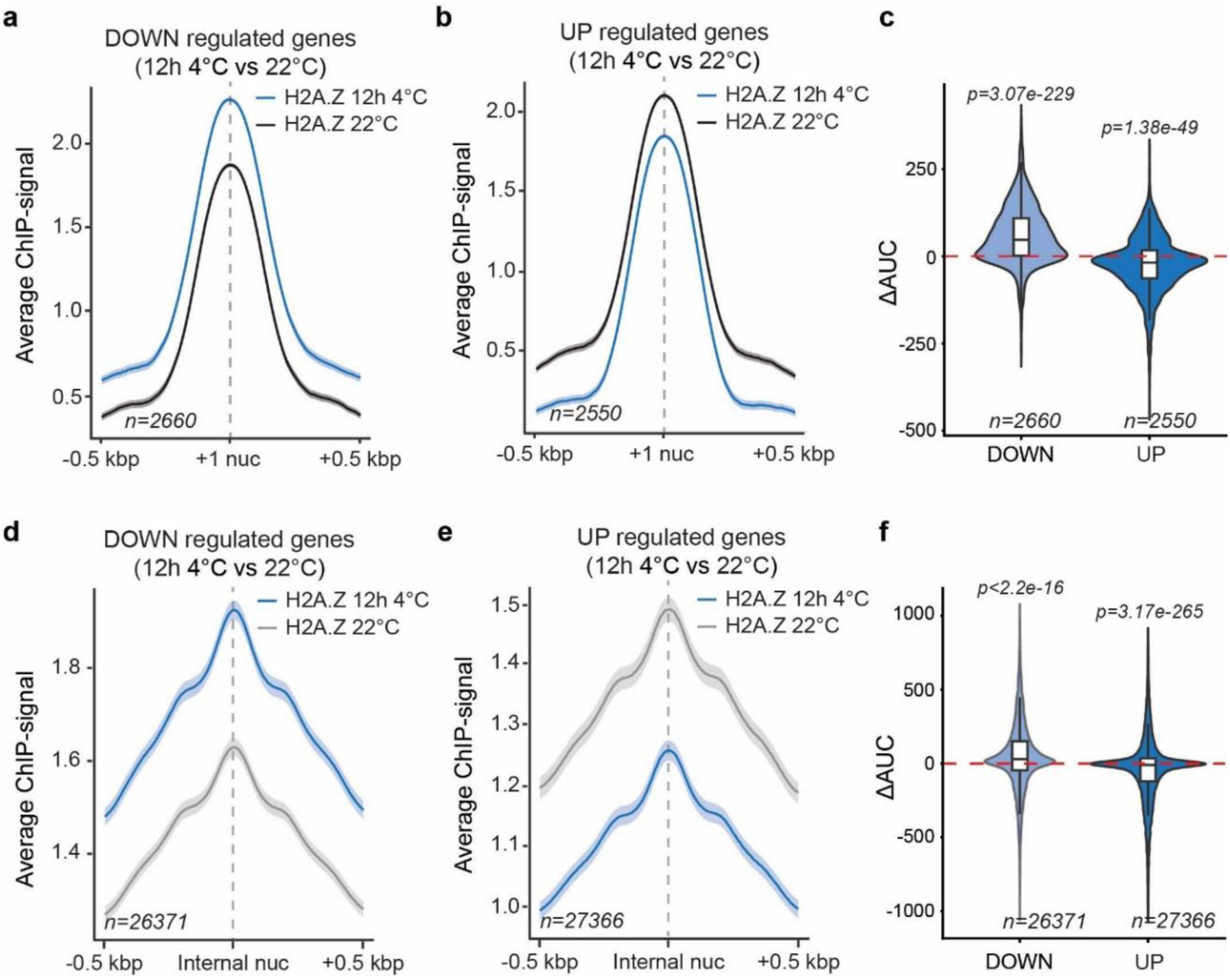
H2A.Z levels change at both the +1 and internal nucleosomes in response to low temperature. a-b, d-e) Metagene plots of average signal from ChIP-seq of H2A.Z at 22°C (in black or in grey) or 12h 4°C (in blue) of down-regulated **(a, d)** or up-regulated genes **(b, e)** at 12h in cold with common peaks of expressed genes in cold at the +1 nucleosome (first nucleosome downstream the TSS) **(a-b)** or internal nucleosomes (nucleosomes under the peak in the gene body without +1 nucleosome) **(d-e)**. The metagene plot is centered on the nucleosome center and includes 500 bp flanks upstream and downstream of the nucleosome center (+1 or internal). The shaded area shows a 95% confidence interval for the mean. **c, f)** Quantification of the metagene plots based on the difference between the areas under the curves (AUC) at 12h 4 °C compared to 22 °C for H2A.Z at the +1 nucleosome (**c**) or at the internal nucleosomes (**f**) or for UP and DOWN gene sets. Positive ΔAUC values indicate higher H2A.Z levels after 12 h at 4 °C compared to 22 °C. The red dashed line marks ΔAUC = 0 (no change). Statistical significance of differences was assessed using two-sided Wilcoxon signed-rank tests to determine whether the median ΔAUC was significantly different from zero, p < 0.05 was considered significant.

**Figure 5.**
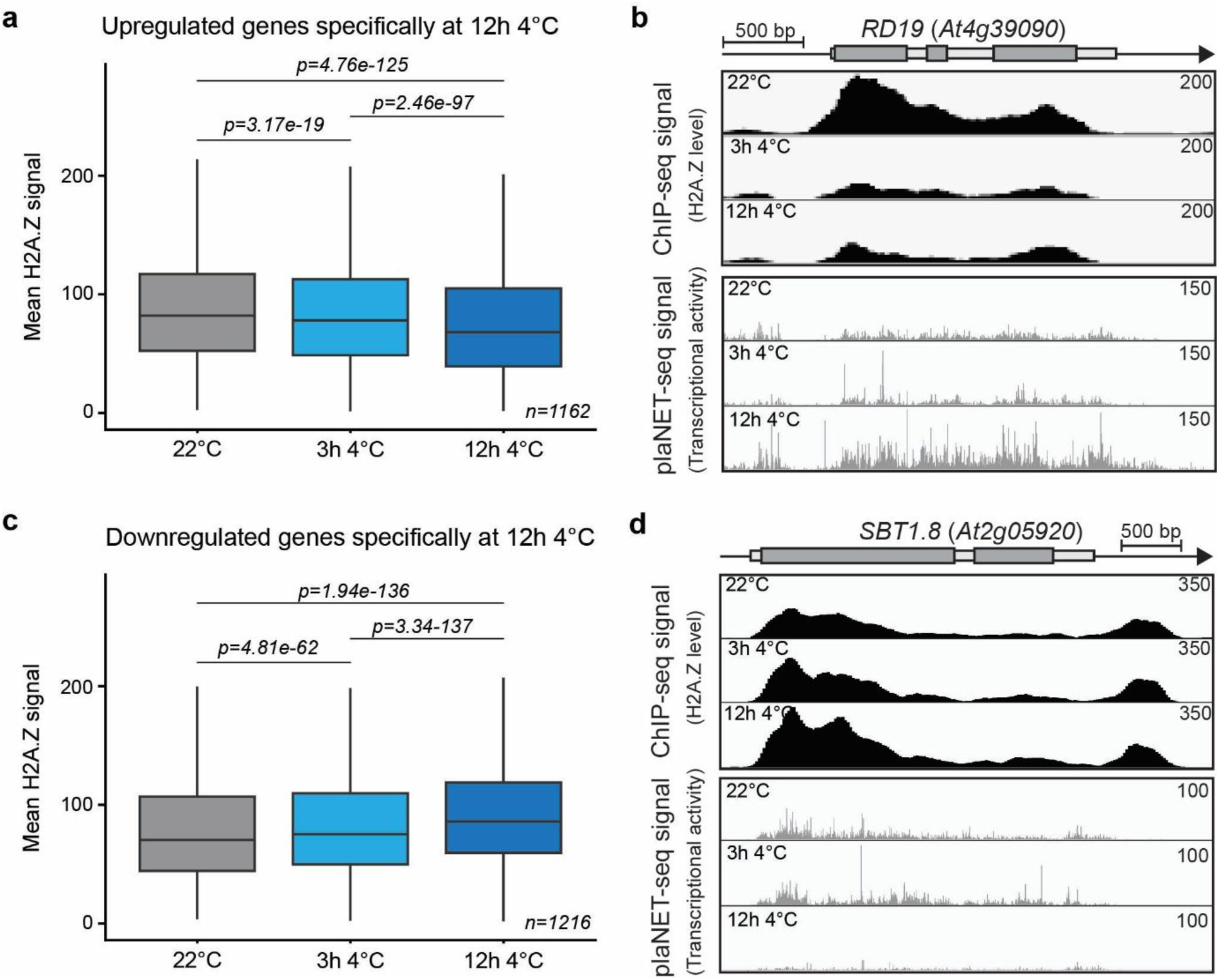
H2A.Z levels change prior to transcriptional changes. a, **c)** Boxplots of H2A.Z enrichment along gene bodies for genes specifically upregulated (**a**) or downregulated (**c**) after 12h at 4 °C and not deregulated after 3h at 4 °C (FC<1.5). H2A.Z levels, extracted from BigWig coverage tracks, are shown for the three time points: 22 °C (in grey), 3 h 4 °C (in light blue) and 12 h 4 °C (in dark blue). Centre lines show the medians; box limits indicate the 25th and 75th percentiles; whiskers extend 1.5 times the interquartile range from the 25th and 75th percentiles. p-values indicate statistical significance of differences between conditions, assessed using two-sided paired Wilcoxon signed-rank tests, p < 0.05 was considered significant. **b, d)** Examples of specific DEG at 12h 4°C: RD19, At4g39090 for up-regulated genes **(b)** and SBT1.8, At2g05920 for down regulated genes **(d)**. Screenshots are from ChIP-seq of H2A.Z and plaNET-seq datasets. Higher occupancy of H2A.Z and elevated transcriptional activity are indicated by higher peak density and amplitude.

To show a biological role of changing H2A.Z levels in response to cold, we used two mutants with altered H2A.Z levels. The *esd1-10* mutant is a knock-out of *EARLY IN SHORT DAYS 1* and its gene product, ACTIN-RELATED PROTEIN 6 (ARP6), is part of the SWI2/SNF2-Related 1 Chromatin Remodelling Complex (SWR1) that incorporates H2A.Z into nucleosomes^36,37^. *HISTONE H2A PROTEIN 9* and *11* are two of the three H2A.Z encoding genes in the Arabidopsis genome and the *hta9-1hta11-2* double mutant leads to an overall lower level of H2A.Z in the genome^27,38^. We selected 5 cold-induced genes with cold-depleted level of H2A.Z and 5 cold-repressed genes with cold-enriched level of H2A.Z (Supplementary Table 7, examples can be seen in Figure 6a-b) for expression analysis at 22°C and after 12h at 4°C. The H2A.Z profile in *arp6-1* mutant at these genes, based on published data^39^, showed reduced H2A.Z enrichment at 22°C (Supplementary Figure 5). Indeed, we detected a misregulation of both sets of genes in the mutants, upregulated genes exhibit reduced activation, while, surprisingly, downregulated genes display enhanced repression, in contrast to the expected attenuation of downregulation (Figure 6c-e), indicating that the level of H2A.Z has a role for gene regulation in response to cold, although this effect appears more consistent for upregulated genes. Together, these results corroborate the hypothesis that H2A.Z levels may function as a precondition for transcriptional change at a subset of genes. It indicates that the H2A.Z levels along the gene body, not only restricted at the +1 nucleosome, are a strong determinant for the level of transcriptional activity in cold conditions. Moreover, it suggests that H2A.Z is an important cold-induced switch, heralding transcription change, for the transcriptional cold response in Arabidopsis.

**Figure 6.**
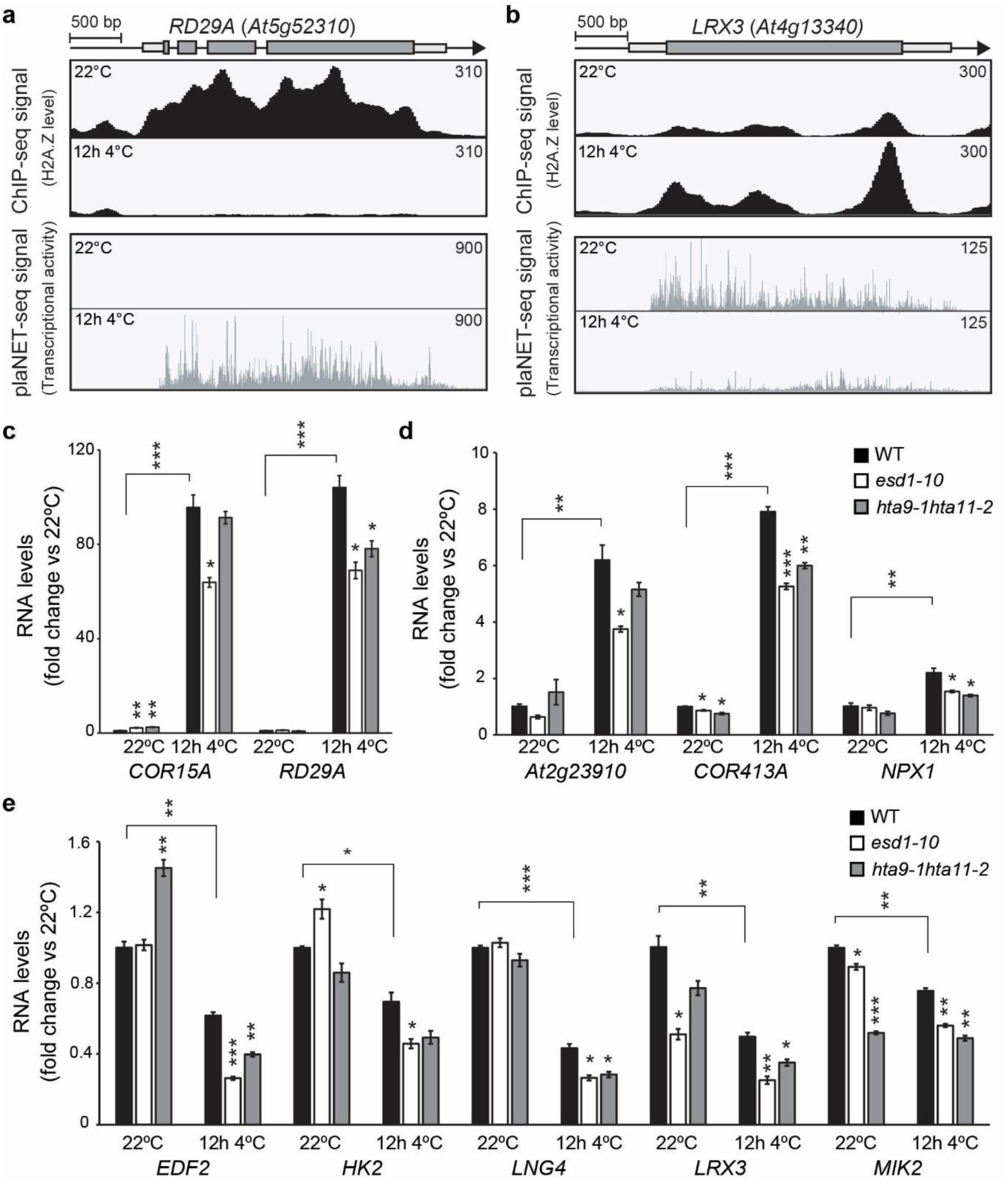
Deposition/Insertion of H2A.Z is required for proper gene regulation in response to low temperature. a-b) Examples of genes with changing H2A.Z levels and regulated in cold (12h 4 °C): RD29A, At5g52310 for up-regulated genes with lower H2A.Z level **(a)** and LRX3, At4g13340 for down regulated genes with higher H2A.Z level **(b)**. Screenshots are from ChIP-seq of H2A.Z and plaNET-seq datasets. Higher occupancy of H2A.Z and elevated transcriptional activity are indicated by higher peak density and amplitude. **c-e)** The relative steady-state level of 5 up-regulated (**c-d**) or down-regulated (**e**) genes regulated in cold (12h 4 °C) with changing H2A.Z levels in WT, esd1-10 and hta9-1hta11-2 measured by RT-qPCR at 22 °C and following 12 h of cold exposure (4 °C). All levels were normalized to WT levels at 22 °C. The mean values are from 3 biological replicates. Error bars represent ± SEM. Statistical significance was calculated with Student’s t-test (* p < 0.05, ** p < 0.01, *** p < 0.001).

### Impaired levels of H3K27me3 result in improper transcriptional cold response

In contrast to H2A.Z levels, H3K27me3 showed no correlation to transcriptional activity of the cold-responsive genes (Figure 2). However, we detected a small but significant depletion of the mark over +1 and internal nucleosomes after 12h at 4°C for differentially expressed genes (Figure 7). The depletion was observed for both down- and up-regulated genes, corroborating that H3K27me3 levels are not correlated with transcriptional activity. In Arabidopsis, there are several H3K27me3 demethylases, including REF6, EARLY FLOWERING 6 (ELF6), and JUMONJI 30 (JMJ30)^40^. Of these REF6 has been found to have a preferred binding site sequence (CTCTGYTY), enabling identification of direct targets^41,42^. Hence, to investigate whether there is a biological effect for the global decrease in H3K27me3 levels, we first identified targets for REF6 among the differentially expressed genes in plaNET-seq after 12h 4°C with a peak of H3K27me3, using REF6 ChIP-seq data^42^. We focused on genes where a REF6 peak was detected in the promoter region of the target gene, representing 70 genes (Supplementary Table 8). Next, we used the *ref6-3* mutant to show the consequences of mis-regulation of demethylation at 5 randomly chosen down- and up-regulated genes from our list. The H3K27me3 profile in *ref6-5* mutant at these genes, based on published data^43^, showed higher H3K27me3 enrichment compared to the WT at 22°C (Supplementary Figure 6). For the up-regulated genes we detected a significant mRNA steady state level change in the wild type seedlings in response to cold (Figure 8a). However, in *ref6-3*, the upregulation was abolished for all investigated genes, indicating that H3K27me3 demethylation is important for the induction of these genes (Figure 8a). Down-regulated genes showed a similar effect. Here, the steady state levels of the target genes in *ref6-3* exhibited increased downregulation compared to wild type (Figure 8b). All in all, these results show that H3K27me3 level control via REF6 is required for the transcriptional response to cold stress. Importantly, H3K27me3 levels do not change in response to transcriptional activity but may rather function as an instructional mark for interacting transcriptional regulators.

**Figure 7.**
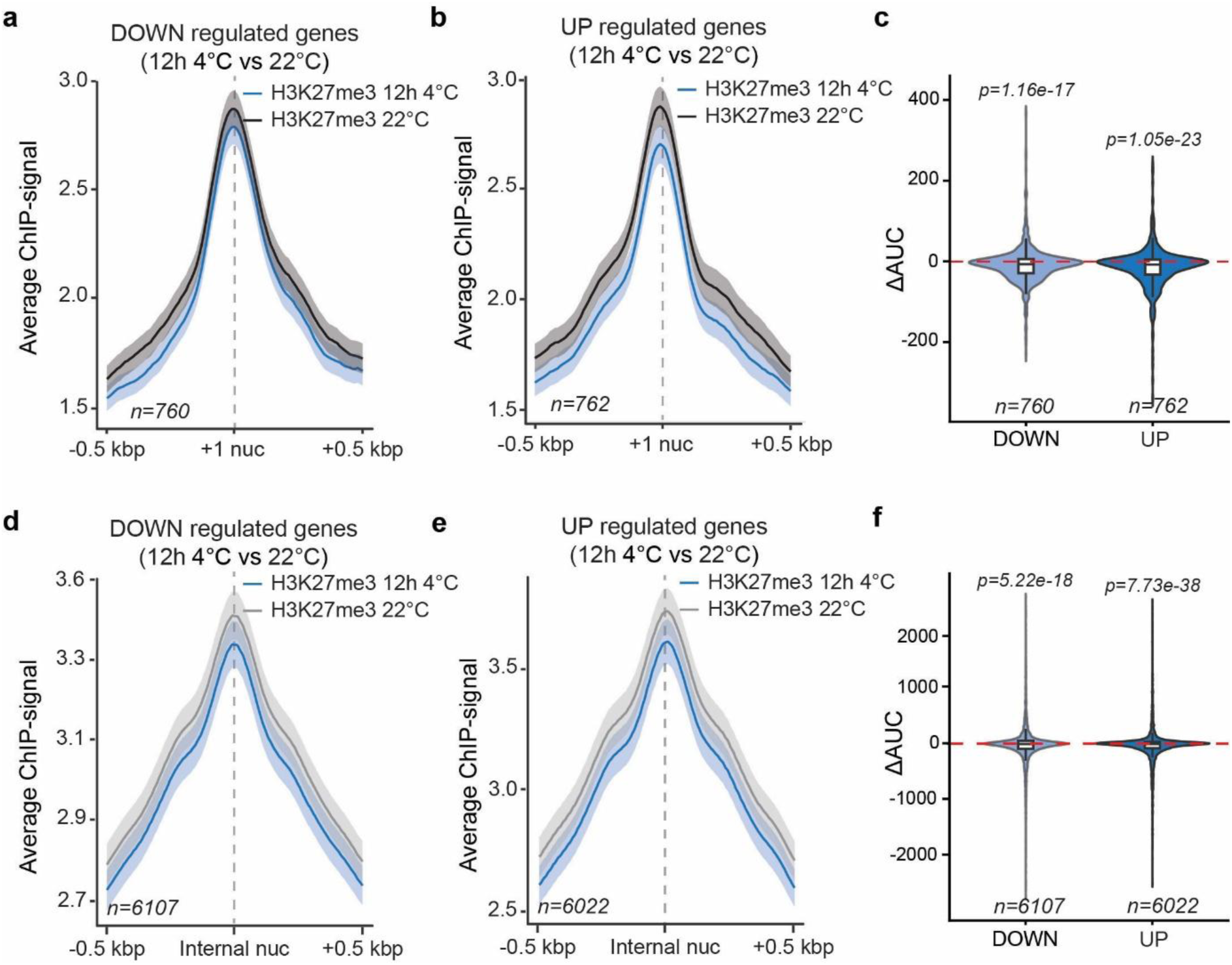
H3K27me3 levels are depleted at both the +1 and internal nucleosomes in response to low temperature. a-b, d-e) Metagene plots of average signal from ChIP-seq of H3K27me3 at 22°C (in black or in grey) or 12h 4°C (in blue) of down-regulated **(a, d)** or up-regulated genes **(b, e)** at 12h in cold with common peaks of expressed genes in cold at the +1 nucleosome (first nucleosome downstream the TSS) **(a-b)** or internal nucleosomes (nucleosomes under the peak in the gene body without +1 nucleosome) **(d-e)**. The metagene plot is centered on the nucleosome center and includes 500 bp flanks upstream and downstream of the nucleosome center (+1 or internal). The shaded area shows a 95% confidence interval for the mean. **c, f)** Quantification of the metagene plots based on the difference between the areas under the curves (AUC) at 12h 4 °C compared to 22 °C for H3K27me3 at the +1 nucleosome (**c**) or at the internal nucleosomes (**f**) or for UP and DOWN gene sets. Positive ΔAUC values indicate higher H3K27me3 levels after 12 h at 4 °C compared to 22 °C. The red dashed line marks ΔAUC = 0 (no change). Statistical significance of differences was assessed using two-sided Wilcoxon signed-rank tests to determine whether the median ΔAUC was significantly different from zero, p < 0.05 was considered significant.

**Figure 8.**
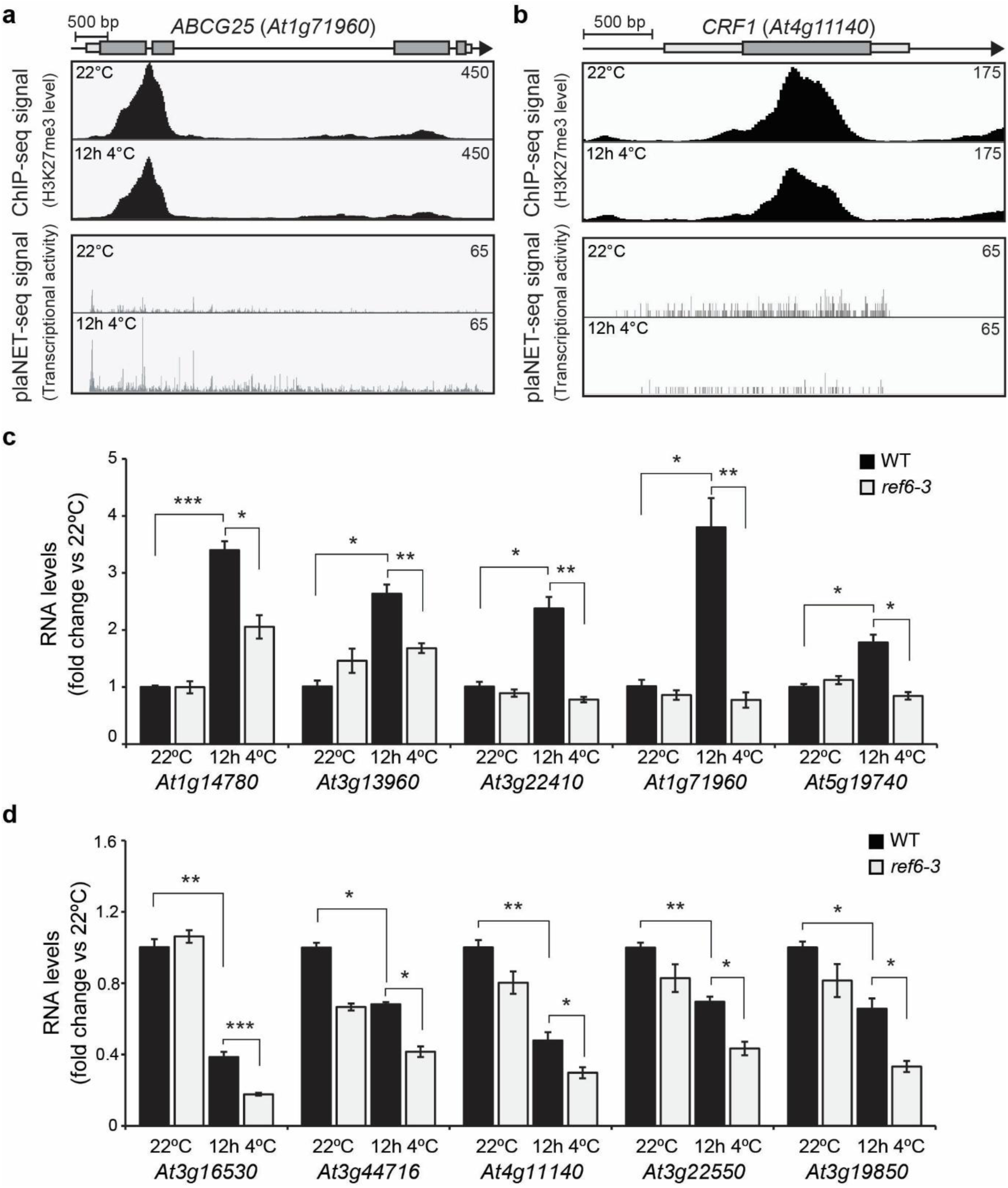
Depletion of H3K27me3 is required for proper gene regulation in response to low temperature. a-b) Examples of target genes of REF6 in their promoter and deregulated in cold (12h 4 °C): ABCG25, At1g71960 for up-regulated genes **(a)** and CRF1, At4g11140 for down regulated genes **(b)**. Screenshots are from ChIP-seq of H3K27me3 and plaNET-seq datasets. Higher occupancy of H3K27me3 and elevated transcriptional activity are indicated by higher peak density and amplitude. **c-d)** The relative steady-state level of 5 up-regulated (**c**) or down-regulated (**d**) genes target of REF6 in their promoter and deregulated in cold (12h 4 °C) in WT and ref6-3 measured by RT-qPCR at 22 °C and following 12 h of cold exposure (4 °C). All levels were normalized to WT levels at 22 °C. The mean values are from 3 biological replicates. Error bars represent ± SEM. Statistical significance was calculated with Student’s t-test (* p < 0.05, ** p < 0.01, *** p < 0.001)

### Transcription over nucleosomes with high or low levels of H2A.Z and H3K27me3 reveal their effect on transcription speed

So far, our results indicated that controlled H2A.Z and H3K27me3 levels are required for plants to respond to cold, albeit with different mechanisms. A potential model would be that changes to marks and variants alter the transcription speed over affected nucleosomes as a mechanism to change transcriptional activity. To understand the direct effects on RNAPII elongation, we classified expressed genes from plaNET-seq data according to relative histone mark signal intensity at each time point (high-signal = top 10% and low-signal = bottom 10%; see Materials and Methods; Supplementary Table 9) and identified +1 and internal nucleosomes (Supplementary Figures 7-8). Then, we calculated the Promoter-proximal Stalling Index (PSI) at the +1 nucleosome (Figure 9a, Supplementary Table 10), from plaNET-seq data, which is a measurement of RNAPII slowing down around the nucleosome^32^, with values around 1 indicating no strong promoter-proximal accumulation and higher values reflecting increased RNAPII pausing. Low H2A.Z levels resulted in decreased promoter stalling compared to enriched levels (Figure 9b), suggesting that high levels of H2A.Z slow down RNAPII. This characteristic was kept after cold treatment (Supplementary Figure 9a-b). In contrast, H3K27me3 levels had no, or small effects on the promoter stalling at 22°C nor after cold treatment (Figure 9c, Supplementary Figure 9c-d), reinforcing the hypothesis that H3K27me3 does not directly affect transcription speed.

**Figure 9.**
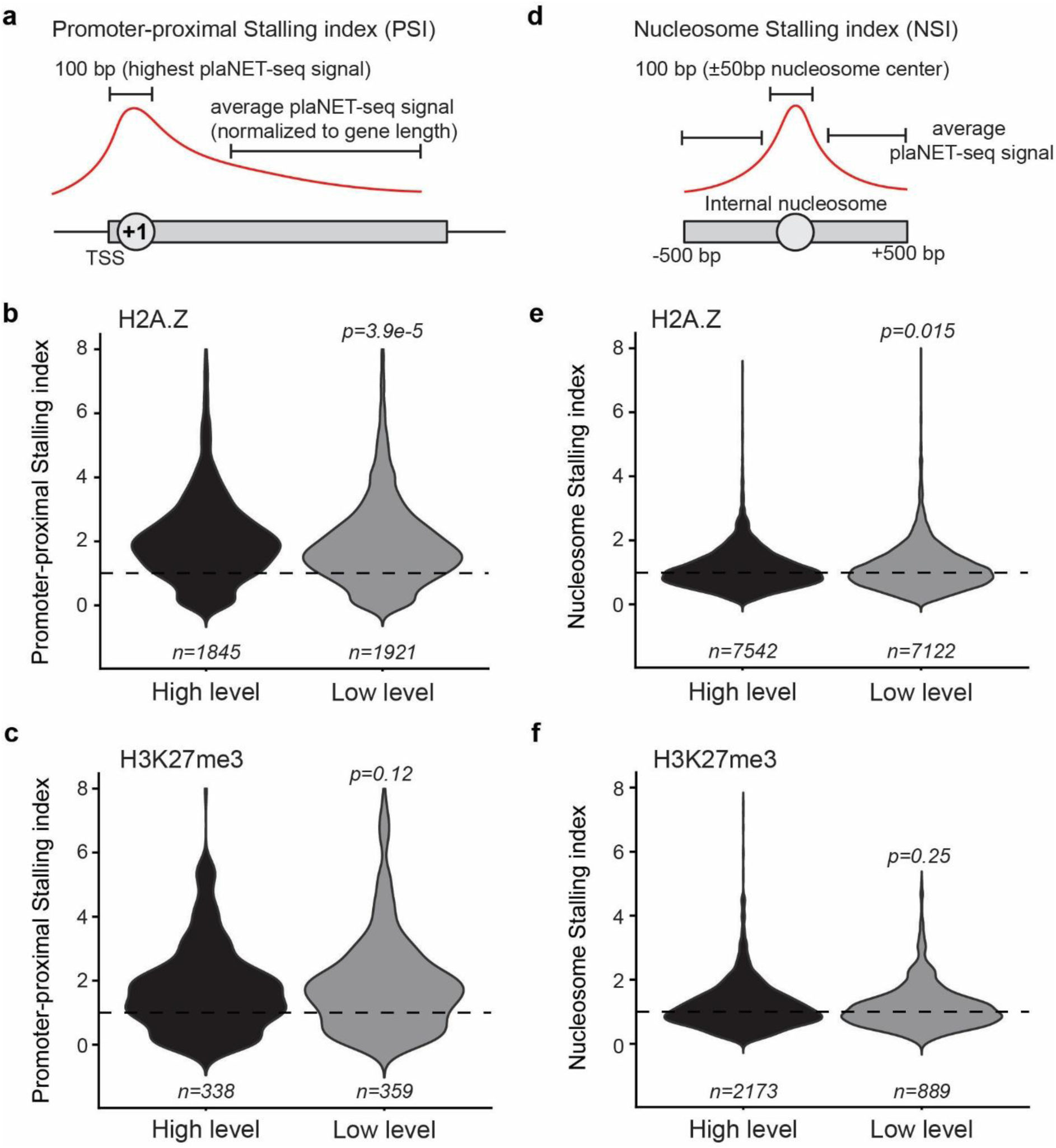
Transcription speed is modulated by H2A.Z but not H3K27me3. a,d) Schematic representation of the calculation of promoter-proximal stalling index (PSI) (**a**) and nucleosome stalling index (NSI) (**d**). **b-c, e-f)** Violin plots showing the promoter-proximal stalling index (PSI) (**b, c**) or the nucleosome stalling index (NSI) (**e, f**) of genes with H2A.Z (**b, e**) or H3K27me3 (**c, f**) peaks at 22 °C. Peaks were classified as high- or low-signal Enriched or Depleted based on levels of the corresponding histone variant or histone modification (10% highest or lowest, respectively). Dashed lines indicate PSI = 1 or NSI = 1, as appropriate. Values around 1 indicate no strong RNAPII accumulation, whereas higher values indicate increased RNAPII pausing. p-values indicate statistical significance of differences between conditions, assessed using two-sided unpaired Wilcoxon signed-rank tests, with p < 0.05 considered significant.

For internal nucleosomes, we calculated a Nucleosome Stalling Index (NSI) by comparing the plaNET-seq signal at the centre of nucleosomes (50 bp up- and downstream of the nucleosome, Supplementary Figure 10a-b, Supplementary Table 11) and the signal surrounding that interval (Figure 9d). Similarly to the +1 nucleosome, genes with low H2A.Z levels showed lower NSI compared to high H2A.Z levels at 22°C and 12h 4°C, but not at 3h 4°C (Figure 9e, Supplementary Figure 9c-d). The difference in NSI was smaller than for PSI, indicating that the levels of H2A.Z have predominantly an effect on the transcription speed over the +1 nucleosome. We could not detect any effect of H3K27me3 levels on transcription speed throughout the time series (Figure 9f, Supplementary Figure 10e-f). Our data reveal that levels of H2A.Z have a direct effect on transcription speed over nucleosomes, but that the position of the nucleosomes impacts the effect more than the variant itself. Moreover, H3K27me3 levels do not have a direct effect on transcription speed.

Taken together, our study highlights the importance of nucleosome content and modification to acclimate the genome to low temperatures and offers a more nuanced view of how histone variants and marks are perceived and regulated by plants.

## DISCUSSION

In this study, we have combined nascent transcription dynamics with epigenomic profiling of H2A.Z and H3K27me3 during short-term cold exposure in Arabidopsis seedlings (3h and 12h at 4°C). Our approach benefits from a direct measurement of transcription activity (plaNET-seq) coupled with changing histone variant composition or H3 modifications (ChIP-seq) in contrast to the more common method of using steady state levels of mRNA (RNA-seq) to describe expression levels. Our study reveals a reorganization of the chromatin environment in response to cold. However, a key point is that a changing chromatin environment is not strictly correlated to transcription activity, rather, it is an acclimatization mechanism to cold temperatures. Furthermore, histone marks and variants described as activating and repressing need to be seen in a much more nuanced view, depending on the level, the position in the gene body, and time of cold exposure.

### Changes in H3K27me3 levels do not directly regulate transcriptional activity

We show that levels of H3K27me3 have small or no direct effects on RNAPII dynamics (i.e. stalling patterns), albeit there is a clear global correlation between transcriptional activity and H3K27me3 levels at 22°C. However, at cold-responsive genes both at control conditions and generally after cold exposure this correlation disappears as the cold-responsive genes show high H3K27me3 enrichment independently of their expression level, questioning a purely “repressive” role of H3K27me3 and suggesting that these genes may adopt more complex chromatin states, potentially involving combinations with other modifications. While transcriptional repression by H3K27me3 via PRC2 deposition is well established in many eukaryotes^44–46^, our results point to a wider role for H3K27me3 in the cold response of plants. We detected numerous genes with changing H3K27me3 level in response to cold, indicating that it may be part of the cold acclimation process. H3K27me3 is believed to increase chromatin compaction in human stem cells^47^, but data from Arabidopsis exposed to low temperature promotes a more nuanced function for H3K27me3^11,12^. Our results indicate that cold temperature alters the distribution of H3K27me3, a mark that clusters nucleosomes and contributes to short- and long-range chromatin interactions.

This seems to be a result of highly dynamic local genome organization. High-resolution mapping of chromatin interactions by chemical cross-linking-assisted proximity capture (CAP-C) revealed that global chromatin conformation remains relatively stable during cold exposure^48^. Our data indicates that the depletion of H3K27me3 and therefore lower chromatin compaction at facultative heterochromatin is largely independent of transcriptional activity. Corroborating this hypothesis, several studies have found that some, but not all, induced genes decrease their H3K27me3 levels in response to cold^1,6,7^. Furthermore, we found that both up- and down-regulated genes are mis-regulated in a *ref6* mutant, a major H3K27me3 demethylase in Arabidopsis. In contrast to cold, warm temperatures lead to a VIN3-LIKE 1 (VIL1)-dependent accumulation of H3K27me3^49^, suggesting that H3K27me3 is indeed a way for the plant to regulate chromatin compaction in response to temperature. Thus, the consequence of regulating H3K27me3 may be to enable other regulatory factors to determine transcriptional activity, not the other way around. The different functional role and localization of H3K27me3 in plants along the gene body compared to animals presumably makes this mechanism plant-specific^10^. In conclusion, we propose that H3K27me3 is not strictly a repressive histone mark, but an important DNA accessibility factor that is required for proper regulation of transcriptional activity after exposure to changing temperatures.

### The H2A.Z levels dictate transcriptional activity during cold exposure

H2A.Z is mainly a conserved feature in most eukaryotes at the +1 nucleosome^50–52^. The +1 nucleosome is well positioned downstream of the promoter to determine transcription start site for RNAPII in eukaryotes^53,54^. In expressed genes, H2A.Z accumulates at this nucleosome^26,55^. Furthermore, it is well established that H2A.Z serves important functions at this nucleosome and post-translational modifications of H2A.Z are linked to gene activation (acetylation)^29^, and repression (mono-ubiquitination)^27^. However, our analysis in Arabidopsis showed that changes to H2A.Z levels (during cold exposure) are largely due to changes in differentially expressed genes along the whole gene body. Furthermore, genes that will be differentially expressed after cold exposure (i.e. cold-responsive genes) have increased H2A.Z levels along the gene body even before cold exposure, compared to non-differentially expressed genes. The correlation between transcription activity and level of H2A.Z suggests that it is more likely to be part of the transcriptional cold response rather than a genome adaptation. In our data, we show that H2A.Z levels at cold-responsive genes, without assessing its potential post-translational modifications, do not correlate with transcriptional activity at 22°C. However, after cold exposure, the H2A.Z levels at +1 and internal nucleosomes, are modified according to increased or decreased transcriptional activity and thus correlate well with transcriptional activity. Importantly, changes in H2A.Z levels precede the transcription change, at least for a subset of genes, suggesting that H2A.Z levels are modified prior, not as a response to transcriptional adaptation. Several lines of evidence support this. Firstly, incorporation of H2A.Z at the +1 nucleosome has been found to lower the energy barrier for DNA wrapping which leads to spontaneous DNA unwrapping^25^, suggesting that incorporation of H2A.Z is important for transcription regulation. Secondly, such spontaneous unwrapping may expose important initiation sequences as in *FLOWERING LOCUS C* (*FLC*)^56^. At this locus, cold temperature-induced removal of H2A.Z leads to repression, indicating that the incorporation of H2A.Z must precede transcriptional regulation. Thirdly, H2A.Z eviction has been found to be transcription independent^18^. These results highlight a conundrum. How can the H2A.Z level of nucleosomes that are more dynamic and easier to unwrap increase in cold-repressed genes and decrease in cold-induced genes at nucleosomes? Our mutant analysis revealed that, as expected, cold-induced genes showed reduced induction, while cold-repressed genes displayed enhanced repression rather than attenuation. This suggests that H2A.Z does not simply promote transcriptional activation but rather modulates transcriptional responsiveness in a more nuanced manner. One possible explanation is that H2A.Z maintains a poised chromatin state that buffers both activation and repression. Its depletion may lead to exaggerated transcriptional responses by affecting chromatin accessibility and RNAPII dynamics. Stress-responsive genes in Arabidopsis accumulate H2A.Z at normal growing temperatures at the +1 nucleosome^30^, which we show correlates with increased RNAPII stalling, a hallmark for stress responsiveness in Arabidopsis^33^. High H2A.Z levels also lead to a slower rate of RNAPII pause release in mouse embryonic cells^57^, indicating that, although an H2A.Z containing nucleosome is easier to unwrap, it also, somewhat counterintuitive, leads to increased RNAPII stalling early in the transcription cycle and during elongation. Additional evidence suggests that the unwrapping of DNA from H2A.Z containing nucleosomes is temperature dependent, with DNA more tightly wrapped at colder temperatures^18^. In fact, both plants and yeast may utilize H2A.Z levels at the +1 nucleosome as thermosensors^18^. All in all, this points to a key active role for H2A.Z in plant transcriptional response to different temperatures and provides an elegant solution for adapting transcription when biochemical process kinetics change^32^.

### The decoration of internal nucleosomes and transcription speed

A kinetic model for histone modification would predict that transcription slows down or picks up speed when H2A.Z or H3K27me3 levels change at internal nucleosomes of expressed genes. We detect small, but significant, differences in RNAPII stalling at nucleosomes with depleted or enriched levels of H2A.Z. However, the effect on RNAPII stalling is larger at the +1 nucleosome where increased stalling is observed at H2A.Z-enriched nucleosomes. Overall, this suggests that indeed H2A.Z containing nucleosomes can affect transcription speed. The differences in RNAPII stalling between +1 and internal nucleosomes make a kinetic model unlikely for H2A.Z, other interacting components are probably required to fully explain the changes to RNAPII dynamics. In fact, H2A.Z has been found to interact with several proteins involved in transcriptional regulation^58^. Further, our results do not support a kinetic model for H3K27me3, as we could not detect any difference in RNAPII dynamics over +1 nucleosomes with enriched or depleted H3K27me3 levels, and we detected no, or small, differences for internal nucleosomes with distinct H3K27me3 levels. This indicates that the H3K27me3 modification is mainly recognized by interacting players to regulate transcription^59^. For example, REF6, a major H3K27me3 demethylase in plants, interacts with a specific motif in the promoter and can recruit light regulated transcription factors^60^, and has been shown to be required for gene activation during germination by promoting an H3K27me3-depleted chromatin state associated with cell identity transitions^61^. A caveat here is that we assume that the investigated marks and variants are not combined with other nucleosome modifications. Thus, an interesting avenue for future research is to combine a range of modifications in samples from similarly aged plants and conditions to fully understand their role in transcription regulation.

Taken together, our study provides important findings of how plants cope with cold temperatures. When temperatures plunge, plants respond by changing its chromatin environment, both to directly regulate transcription and adapt the chromatin to the new condition. It highlights that the composition and modification of nucleosomes have a more nuanced influence on transcription than previously believed, especially if the plants are grown under control conditions and compared to a stress condition.

## MATERIALS AND METHODS

### Plant growth and conditions

Wild-type Arabidopsis (*Arabidopsis thaliana*) ecotype Columbia-0 (Col-0) and the T-DNA insertion mutants *ref6-3* (SAIL_747_A07)^62^, *arp6/esd1-10* (WiscDsLox289_292L8)^63^ and *hta9-1 hta11-2* (SALK_054814; SALK_031471)^38^ were used in this study. Seeds were surface sterilized and stratified for 2 days at 4°C in the dark. Seedlings were grown on ½ Murashige and Skoog (MS) basal medium supplemented with 1% (w/v), with long day conditions (16h light at 22°C, 8h dark at 18°C, ∼100 µEm^−2^ s^−1^) for 10 days in SciWhite LEDs (Percival) for RT-qPCR or in Aralab growth chamber for ChIP-seq. Cold treatments were performed as described in Kindgren et al.,2020^32^. Briefly, 10-day-old seedlings were transferred from 22°C to 4°C and exposed to ∼25 µEm^−2^ s^−1^ 4 hours after the light onset (ZT4) for 3h or 12h.

### ChIP-seq

ChIP experiments were conducted in two biological replicates for each timepoint. Batches of 1g of 10-day-old seedlings (roots and aerial parts) were harvested for each time point and immediately cross-linked for 15 min using 1% formaldehyde in cold UltraPure^TM^ Distilled Water (Invitrogen) under vacuum with a wire mesh to keep all the seedlings submerged. The crosslink reaction was stopped by the addition 2 M glycine to a final concentration of 0.125 M for 5 min under vacuum. Samples were rinsed twice with cold water, dried briefly on tissue paper and frozen in liquid nitrogen. For each biological replicate, 4 batches of 1g of cross-linked tissues were ground to fine powder using pre-cooled mortar and pestle in liquid nitrogen. Each batch was processed separately for the nuclei extraction steps, then merged by two for the nucleus lysis and sonication, and finally all merged after the sonication to process IP. The nuclei extraction was carried out as follows, ground tissues were put in a cold falcon tube with 25 ml of Extraction buffer 1 (0.4 M sucrose, 10 mM Tris-HCl pH 8, 10 mM MgCl_2_, 5 mM beta-mercaptoethanol, 0.1mM phenylmethylsulfonyl fluoride [PMSF], 1X cOmplete protease inhibitor cocktail (Roche)) were added to the powder, samples were vortexed. The solution was filtered through 1 layer of miracloth (VWR) and centrifuged for 20 min at 4000 rpm at 4°C. Supernatant was removed and pellet was washed at least four times (1:1ml, 2-3: 500µl, 4: 1ml) until pellet was not green anymore with Extraction buffer 2 (0.25 M sucrose, 10 mM Tris-HCl pH 8, 10 mM MgCl_2_, 1% Triton X-100, 5 mM beta-mercaptoethanol, 0.1mM PMSF, 1X cOmplete protease inhibitor cocktail (Roche)). Samples were resuspended in 300 μl Extraction buffer 3 (1.7 M sucrose, 10 mM Tris-HCl pH 8, 2 mM MgCl_2_, 0.15% Triton X-100, 5 mM beta-mercaptoethanol, 0.1mM PMSF, 1X cOmplete protease inhibitor cocktail (Roche)) and layered on 300 μl Extraction buffer 3 (sucrose gradient), centrifuged for 1 h at 16,000 g at 4°C. The pellet was resuspended in 105 μl Nuclear Lysis Buffer 0.5 % SDS (50 mM Tris-HCl pH 8, 10 mM EDTA, 0.5% SDS, 0,1mM PMSF, 1X cOmplete protease inhibitor cocktail (Roche)) and pooled per two and then incubated 30 min at 4°C. 840µl of Nuclei Lysis Buffer 0% SDS (50 mM Tris-HCl pH 8, 10 mM EDTA, 0.1mM PMSF, 1X cOmplete protease inhibitor cocktail (Roche)) were added to the samples to reach a final concentration of SDS at 0.1 % and a volume of 1.05mL for the sonication and incubated 45 min at 4°C with rotation. Chromatin was sonicated using an S220 Focused-Ultrasonicator (Covaris) with the following settings: treatment time 5 min, acoustic duty factor 20%, PIP 175W, Cycles per burst 200 and temperature 2-5°C to get fragment sizes between 100 and 500 bp. Nuclear debris were removed by centrifugation at 5000 rpm for 10 min at 4°C and kept at 4°C while sonication efficiency and samples concentration were verified. Nuclear debris were removed again by centrifugation at 12000 g for 10 min at 4°C and pooled the two chromatin samples. The same chromatin was used for the different antibodies for each replicate and each timepoint and to always use the same amount of chromatin in the same final volume (chromatin + 0.5 volume Covaris dilution Buffer). If needed, chromatin was diluted to obtain the desired quantity of chromatin with Nuclear Lysis Buffer 0.1 % SDS (50 mM Tris-HCl pH 8, 10 mM EDTA, 0.1% SDS, 0,1mM PMSF, 1X cOmplete protease inhibitor cocktail (Roche)) to adjust concentration of NaCl to 150mM and Triton X-100 to 1% for the immunoprecipitation 0.5 volume of Covaris dilution Buffer (20mM Tris HCL pH8, 2mM EDTA, 0.1%SDS, 450mM NaCl, 3% Triton X-100) was added and one tube per IP was thus prepared. Dynabeads Protein A (Invitrogen - Thermo Fisher Scientific) were washed 3 times with ChIP dilution buffer (16.7 mM Tris pH8, 1.2 mM EDTA, 167 mM NaCl, 1.1% Triton X-100, 0.1%SDS, 0.1mM PMSF, 1X cOmplete protease inhibitor cocktail (Roche)) and resuspended in the initial volume with ChIP dilution buffer then 15 µl of washed beads were added to each chromatin tube and incubated 3h at 4°C with rotation for pre-clearing. Then the samples were quickly spun, the beads were removed and 690µl of supernatant from each was taken to perform IP and the remaining 110 µl of supernatant from each was taken and combined. From this tube, 2 x 20 µl were taken to make two input tubes and the remaining 290 µl was used as a Mock. Inputs were stored at -20°C until the reverse-crosslink step. Immunoprecipitation was performed by adding the appropriate amount of antibody to each IP tube to have a ratio 1µg of chromatin:1µg of antibody; we performed IP with 2 µg of chromatin with antibodies recognizing H3K27me3 (Diagenode C15410069, lot A1818P) and H2A.Z (HTA9; Agrisera, AS10718, lot 2011) and incubated O/N at 4°C with rotation. ChIP dilution buffer from unused beads was removed and replaced by 1 ml of Pre-coating buffer (10µl of Glycogen 20 mg/ml, 10µl of BSA 20 mg/ml, 10µl of yeast t-RNA 20 mg/ml in ChIP dilution buffer) and incubated O/N at 4°C with rotation. Beads were washed 3 times with ChIP dilution buffer and resuspended in the initial volume without volume used for pre-clearing with ChIP dilution buffer. 50μL of prewashed beads were added to the IP and 10μl to the Mock and incubated 3h at 4°C with rotation. The tubes were quickly centrifuged and the beads were pelleted. Beads were washed twice (first a quick wash by vortexing and then a second wash for 5 min at 4°C with rotation) at 4°C in cold-room with Low Salt wash buffer (150 mM NaCl, 0.1% SDS, 1% Triton X-100, 2 mM EDTA, 20 mM Tris-HCl pH 8), High Salt wash buffer (500 mM NaCl, 0.1% SDS, 1% Triton X-100, 2 mM EDTA, 20 mM Tris-HCl pH 8), LiCl wash buffer (0.25 mM LiCl, 1% Nonidet P-40, 1% sodium deoxycholate, 1 mM EDTA, 10 mM Tris-HCl pH 8) and twice quick washes (beads were transferred in a new tubes after first wash) with TE buffer (10 mM Tris-HCl pH8, 1 mM EDTA). Elution was achieved by adding 250 μl of 65°C warm Elution buffer (1% SDS, 0.1 M NaHCO_3_) to the samples and incubating them for 15 min at 65°C with gentle shaking. The elution was repeated and the two eluates of each sample were combined into a single tube and 480 µl of Elution buffer were added to the Input samples to bring the volume to 500µl. The eluates were reverse-crosslinked by adding 20 μl of 5 M NaCl and incubated O/N at 65°C with shaking. Proteins were removed by adding 1 μl of Proteinase K (20 mg/ml, Thermo Fisher Scientific), 10 μl of 0.5 M EDTA and 20 μl of 1 M Tris-HCl (pH 6.5) and incubated for 1 hour at 45°C. Then, samples were treated with 10 μg of RnaseA and incubated for 30min at room temperature. Reverse crosslinked DNA was purified using ChIP DNA clean and concentrator kit (Zymo Research) following the manufacturer’s instructions and eluted in a final volume of 20 μl. Library preparation from the eluted DNA and PE100 sequencing on a DNBseq sequencing platform was performed at the Beijing Genomics institute (Hong Kong).

### The ChIP-seq data processing

The ChIP-seq data were uploaded to the Galaxy web platform, and we used the public server at usegalaxy.org^64^ to analyse the data until alignment and switched from the MarkDuplicates step to usegalaxy.eu^65^. Reads were trimmed and filtered for quality and adapter contamination using Trim Galore! (galaxy version 0.6.7+galaxy1) (https://github.com/FelixKrueger/TrimGalore) and aligned to the TAIR10 genome with Bowtie2^66^ (galaxy version 2.5.3+galaxy1) using the --very sensitive setting. Multi-mapping reads were discarded, and PCR duplicates were removed using Picard Tools’ MarkDuplicates (galaxy version 3.1.1.0) using the remove duplicates option (http://broadinstitute.github.io/picard/). Peaks of H3K27me3 and H2A.Z were called using MACS2^67^ (galaxy version 2.2.9.1+galaxy0) with the following settings: input used as control file, genome size 120e6 (excluded chrM and chrC), no model, extension size of 100, no shift, peaks detection based on q-value, q-value < 0.01, broad regions, broad region cutoff 0.1. Differential enriched peaks between timepoint were defined using DiffBind^68^ (galaxy version 3.12.0+galaxy0) with Input files as control files, DESeq2, only include peaks in at least 2 peak sets in the main binding matrix, FDR<0.05 as settings. Called peaks and DiffBind results were annotated with annotatePeaks^69^ (galaxy version 4.11+galaxy3) using araport11 annotation. Coverage Bigwig and bedgraph files were generated with Bedtools suite tool’s BamCoverage^70^ (galaxy version 3.5.4+galaxy0) using the reads per kilobase per million (RPKM) normalization method for genome browser visualization or Counts per million (CPM) to comparable data for +1 nucleosome and internal nucleosome analysis.

### ChIP-seq analyses and visualization

For intersection analyses of genes associated with differentially enriched peaks, peaks were defined by a fold change > 2 and FDR<0.05. For intersection analyses of peaks with differential signal intensity, peaks were defined as those with an absolute ΔIntensity>1.5× the standard deviation (SD) of all peaks. Mean peak intensities were calculated from the ChIP-seq binding matrix by averaging across replicates for each condition. ΔIntensity was computed as the difference between cold-treated samples (3 h or 12 h at 4°C) and control samples (22°C). The SD was calculated across all peaks within the dataset, and the threshold (1.5× SD) was empirically defined to identify peaks with substantial signal changes. UpSet plots were generated in R with the UpSetR package^71^. Venn diagrams were generated in R using the VennDiagram package^72^ to compare genes associated with differentially enriched peaks (based on fold change) or peaks with differential signal intensity (ΔIntensity) with differentially expressed genes (DEGs) identified by plaNET-seq after cold exposure.

For correlation analyses between ChIP-seq and transcriptional activity, normalized ChIP-seq signals were quantified over extended genic regions (1 kb upstream of the transcription start site plus the gene body) for expressed genes. These signals were correlated with plaNET-seq expression values (log₂(FPKM + 0.1)) using Pearson correlation, reporting the coefficient of determination (*R²*). To assess the relationship between transcriptional response and histone occupancy upon cold exposure, mean peak intensities from the ChIP-seq binding matrix (each replicate and condition) were used to calculate ΔIntensity between cold-treated (3h or 12h at 4°C) and control (22°C) samples. ΔIntensity values were correlated with log₂ fold-changes in plaNET-seq expression using linear regression models for visualization and statistical interpretation.

To classify peaks based on relative signal intensity for each histone mark, high and low level peaks were defined based on absolute ChIP-seq signal intensity at each time point by selecting the 10% highest and 10% lowest of signal intensity, respectively. These categories therefore reflect relative signal levels within a given condition, high- and low-signal peaks, rather than differential enrichment between conditions. Only peaks annotated within genes expressed according to plaNET-seq data were retained, while intergenic or *NA*-annotated regions were excluded. The same peak sets were used for analyses focusing on either the +1 nucleosome or internal nucleosomes.

Metagene plots were generated following the approach described by *Kindgren et al.* (2020)^32^, using the code available on GitHub (https://github.com/Maxim-Ivanov/Kindgren_et_al_2019, 05_Metagenes) with custom adaptations to accommodate internal nucleosome positioning. Metagene plots corresponding to the +1 nucleosome were produced independently using ngs.plot^73^. Quantification was performed by calculating the difference in the area under the curve (ΔAUC) for each gene, defined as (AUC4°C – AUC22°C), where AUC represents the summed signal intensity across the metagene profile.

Promoter-proximal and nucleosome stalling indices (PSI and NSI) were computed as described in *Kindgren et al.* (2020)^32^ (code available on GitHub, 09-Stalling_index), with modifications for the calculation of NSI. NSI values were calculated as the ratio of plaNET-seq coverage within a 100 bp central window (±50 bp) centred on internal nucleosomes under peaks to the mean coverage of two 450 bp flanking regions upstream and downstream. Signals were normalized to window length, and nucleosomes located within 300 bp of either the TSS or the PAS were excluded.

Additional graphical representations, including boxplots and violin plots, were generated using ggplot2 package^74^ in R to visualize signal distributions and variability across experimental conditions.

### RNA extraction, cDNA synthesis and RT-qPCR

The RNA extraction, cDNA synthesis, RT-qPCR and data analysis were performed as described by Rosenkranz et al. 2025^75^. Shortly, 1 µg of total RNA extracted using the Plant RNA Mini Kit (Omega-Biotek) and treated with dsDNase (Thermo Fisher Scientific) was used for complementary DNA (cDNA) synthesis using iScript reverse transcriptase (Bio-Rad). Quantitative real-time PCR (RT-qPCR) was performed on CFX384 Real-Time PCR detection systems (Bio-Rad) using SYBR premix (Bio-Rad). Each time, data from 3 biological replicates (approximately 20-30 seedlings grown on separate plates) with 3 technical replicates were considered. Statistically significant differences were calculated using Student’s t-test. The primers used are listed in the Supplementary Table 12.

### Genome-wide data sets

Available genome-wide datasets used in this study include plaNET-seq (GSE131733)^32^, MNase-seq (PRJA592356)^76^, REF6 ChIP-seq (GSE72736)^42^, RNAPII ChIP-seq (GSE95301)^34^ same analysis as Meena et al., 2024^33^, H3K27me3 ChIP-seq (GSE255444)^12^ H2A.Z ChIP-seq (GSE108450)^35^, H2A.Z ChIP-seq in *arp6-1* mutant (GSE123263)^39^ and H3K27me3 ChIP-seq in *ref6-5* mutant (GSE181292)^43^. Moreover, we used nucleosome occupancy tracks and nucleosome coordinates available from the PlantDHS database (https://plantdhs.org/Download (Nucleosome file from leaf))^77^ for the internal nucleosomes and MNA-seq data (GSE205112)^78^ for the first nucleosome position used in metagene.

## Data availability

ChIP-seq data generated in this study have been deposited in the ENA under BioProject accession number PRJEB108427 including raw data. Bigwig files and peak files are available in SciLifeLab (https://figshare.scilifelab.se/).

## ACKNOWLEDGEMENTS

We would like to thank members of the Kindgren lab for their critical reading of the manuscript. We are grateful to Åsa Strand for the gift of the *ref6-3* mutant line and to Manuel A. Piñeiro for the gift of the *esd1-10* and *hta9-1 hta11-2* mutant lines. We would like to thank the greenhouse personnel at Umeå Plant Science Centre for plant maintenance, the UPSC Bioinformatics facility (https://bioinfomatics.upsc.se) and iGReD Bioinformatics platform (https://bioinformatics-bim.igred.fr) for technical support. We would also like to acknowledge support from Science for Life Laboratory. The research in the P.K. lab was funded by FORMAS (2021-01065), Kempe foundation (JCK-3131), and Novo Nordisk Foundation (NNF24OC0094849). This work was partially supported by the Wallenberg Initiatives in Forest Research (WIFORCE) funded by the Knut and Alice Wallenberg Foundation (2020.0240). We acknowledge support from ANR grants SeedChrom (ANR-22-CE20-0028, to AVP) and EpiLinks (ANR-22-CE20-0001, to SA). AVP and SA thank the GDR EPIPLANT for networking support.

## AUTHOR CONTRIBUTIONS

S.M., S.A., A.V.P and P.K. conceived the study and designed the research. S.M. and P.K. performed the experiments and collected the data. S.M., S.M.N. and P.K. contributed to data analysis and S.M., S.M.N., P.K., S.A. and A.V.P. to interpretation. S.M. and P.K. wrote the original draft of the manuscript. All authors contributed to manuscript revision, read, and approved the final version. P.K. and A.V.P. supervised the project.

## COMPETING INTERESTS

The authors declare no competing interests.

## Notes

### Competing Interest Statement

The authors have declared no competing interest.

