## Supplementary figures for "H3K27me3 and H2A.Z prime cold regulated genes, and their remodelling governs plant cold response"

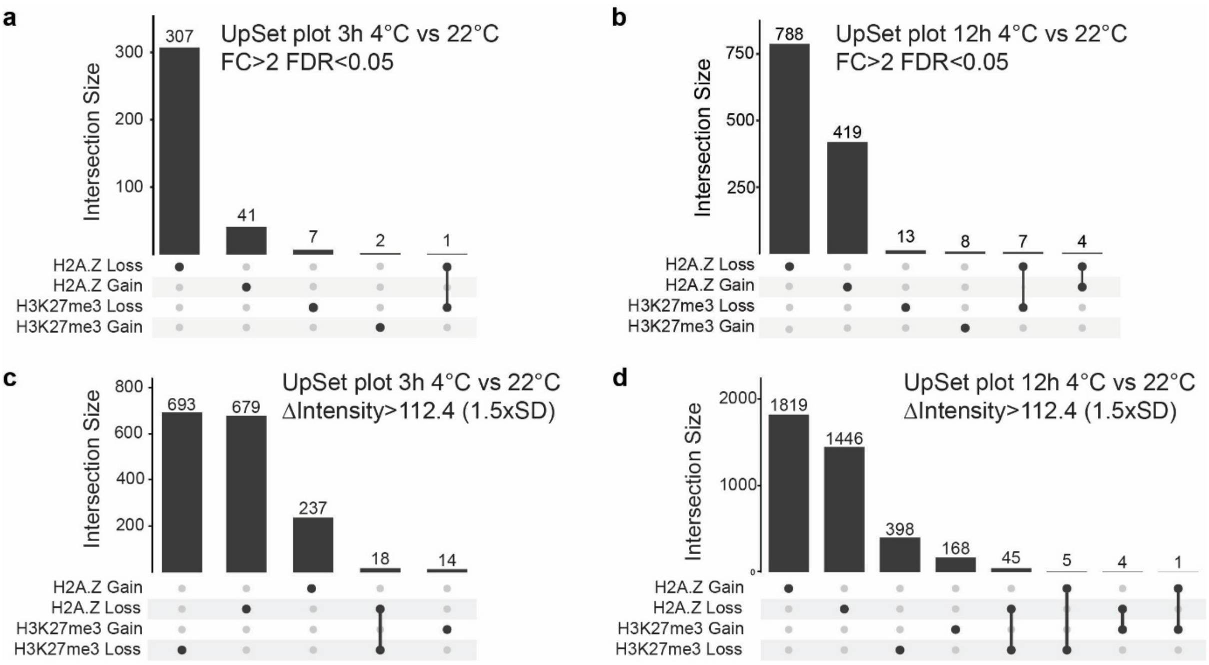


**Supp. Figure 1. UpSet plots showing gene- and peak-level changes in H2A.Z and H3K27me3 in response to cold exposure**

**a-d**) UpSet plots showing intersections of genes associated with differentially enriched peaks (FC>2 and FDR<0.05; **a, b**) or peaks with differential signal intensity (Δintensity>1.5xSD of all peaks; **c, d**) for H2A.Z and H3K27me3 after 3h (**a, c**) or 12h (**b, d**) in cold.

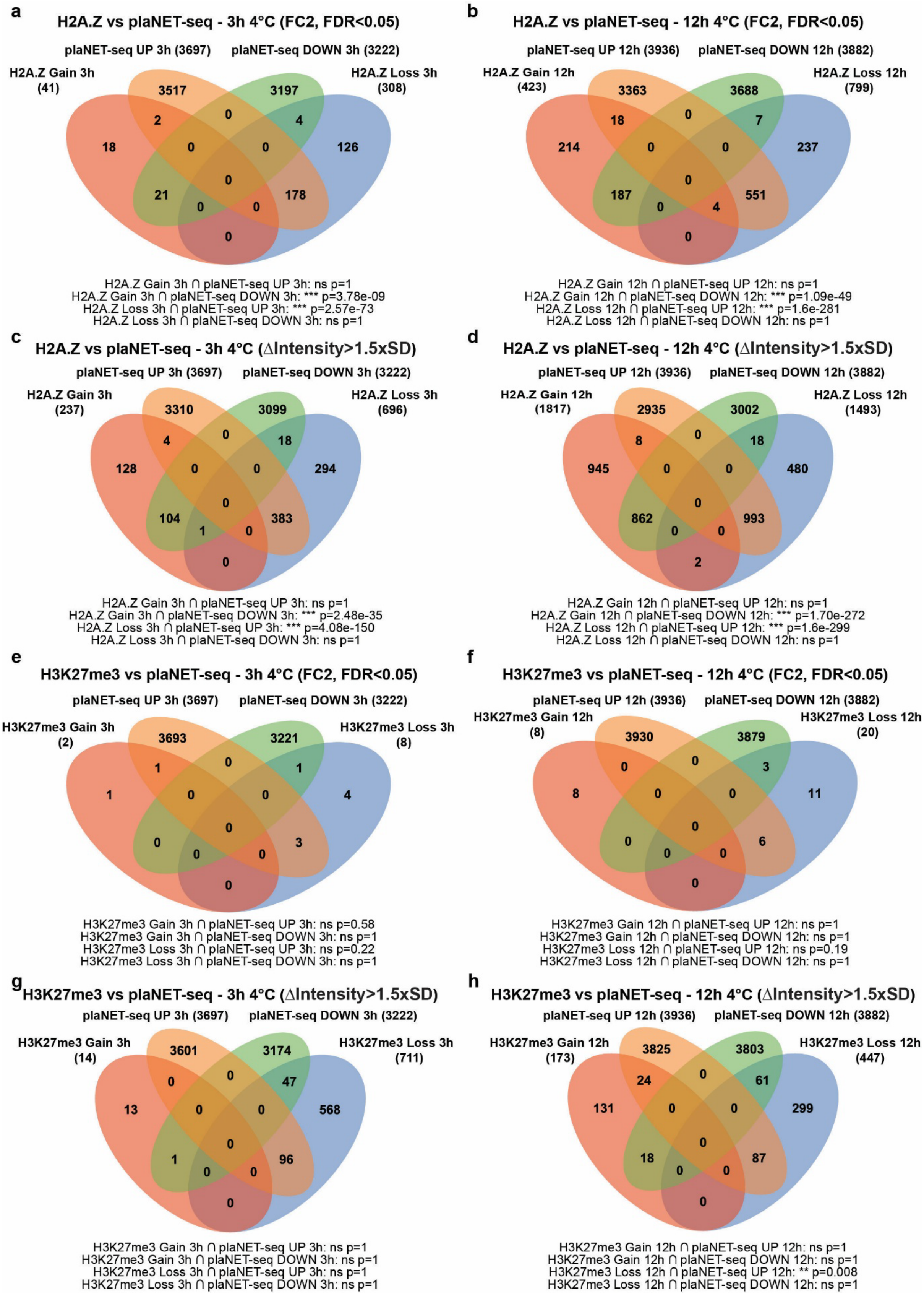


**Supp. Figure 2. Overlap between chromatin and transcriptional changes in response to cold exposure**

**a–d**) Venn diagrams showing the overlap between genes up- or downregulated by plaNET-seq and genes associated with H2A.Z changes (gain or loss), defined either by differential enrichment (FC>2, FDR<0.05; **a, b**) or by differential signal intensity (Δintensity > 1.5×SD of all peaks; **c, d**), after 3 h (**a, c**) and 12 h (**b, d**) of cold exposure at 4°C.

**e–h**) Venn diagrams showing the overlap between genes up- or downregulated by plaNET-seq and genes associated with H2A.Z changes (gain or loss), defined either by differential enrichment (FC>2, FDR<0.05; **e, f**) or by differential signal intensity (Δintensity > 1.5×SD of all peaks; **g, h**), after 3 h (**e, g**) and 12 h (**f, h**) of cold exposure at 4°C.Statistical significance of overlaps was assessed using a one-sided Fisher's exact test based on a hypergeometric distribution. P values were adjusted for multiple testing using the false discovery rate, $p < 0.05$ was considered significant.

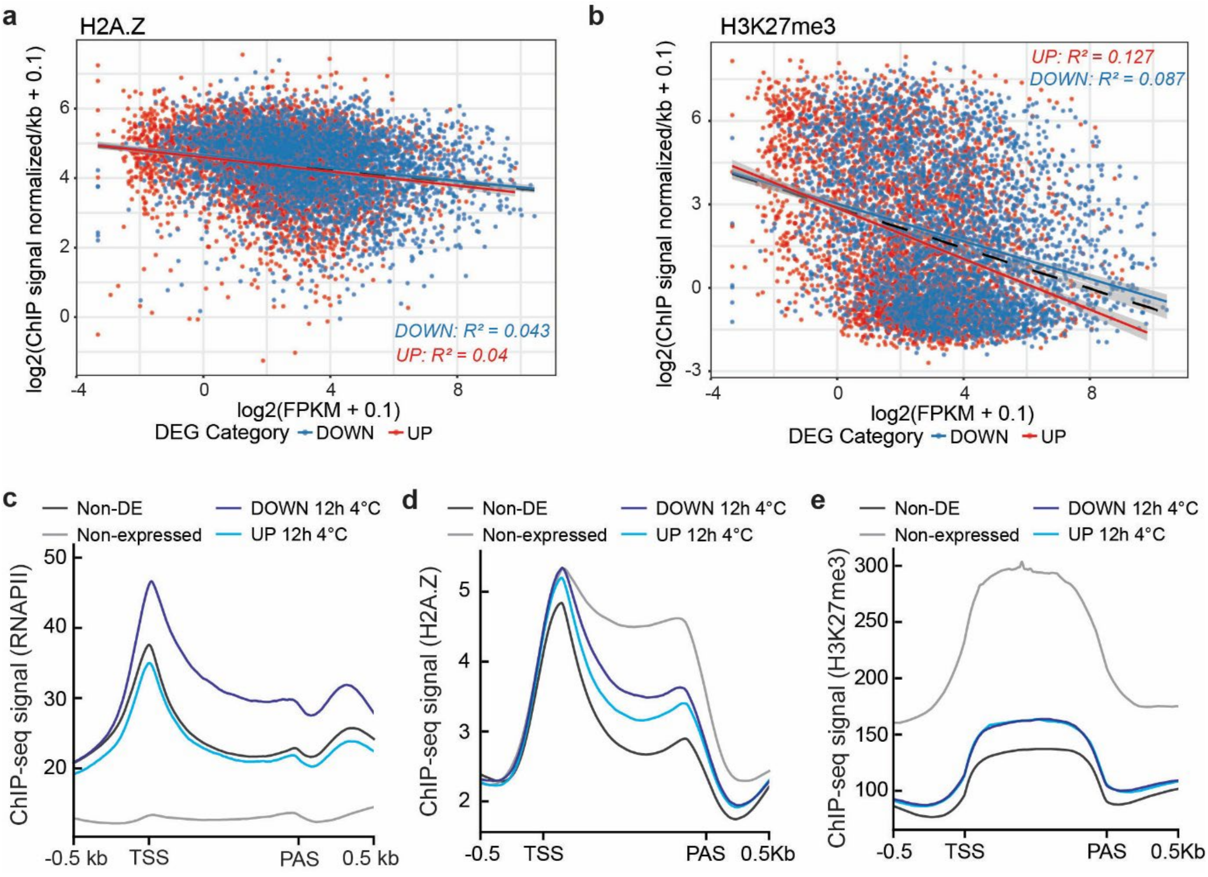


**Supp. Figure 3. Correlations between transcriptional activity and H2A.Z/H3K27me3 levels of genes that are DE after cold treatment at 22°C.**

**a-b)** Correlation plots between ChIP-seq signal and gene expression levels for H2A.Z (**a**) and H3K27me3 (**b**) at 22 °C. Each point represents a gene significantly up regulated (red) or down regulated (blue) after 12 h of cold treatment determined in plaNET-seq. The black dashed line indicates the global linear regression across all genes, while colored lines show regressions for each DEG category. The coefficient of determination ($R^2$) was calculated using Pearson's correlation.

**c-e)** Metagene plots of average signal from published and available data from ChIP-seq of RNA PolII[34] **(c),** H2A.Z[35] **(d)** or H3K27me3[12] **(e)** at 22°C of expressed non-deregulated genes in cold (non-DE, in black), non-expressed genes (in gray), down regulated (DOWN 12h 4°C, in dark blue) or up regulated (UP 12h 4°C, in light blue) genes in plaNET-seq at 12h at 4°C .The metagene plot covers the gene body and includes 500 bp flanking regions upstream and downstream of the TSS and PAS, respectively. TSS: Transcription Start Site, PAS: Poly(A)-Signal.

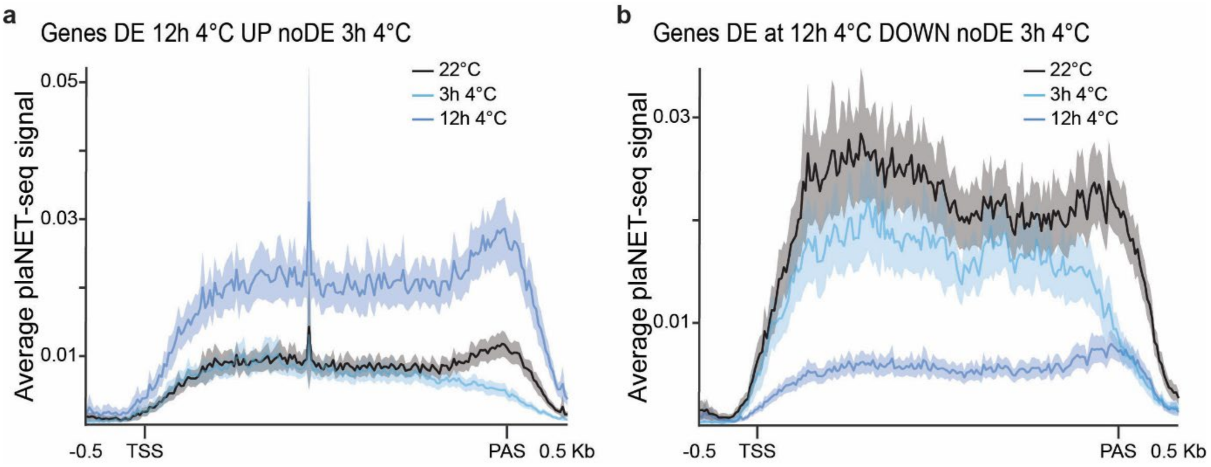


**Supp. Figure 4. Correlations between transcriptional activity and H2A.Z/H3K27me3 levels of genes that are DE after cold treatment at 22°C.**

**a-b)** Metagene plots of average plaNET-seq signal of genes DE UP (**a**) or DOWN (**b**) at 12h 4°C (FC>2) and noDE at 3h 4°C (FC<1.5) (same gene sets as in Fig. 5). Signals at 22°C (black), 3 h at 4°C (light blue), and 12 h at 4°C (dark blue) are indicated. The metagene plot covers the gene body and includes 500 bp flanking regions upstream and downstream of the TSS and PAS, respectively. TSS: Transcription Start Site, PAS: Poly(A)-Signal.

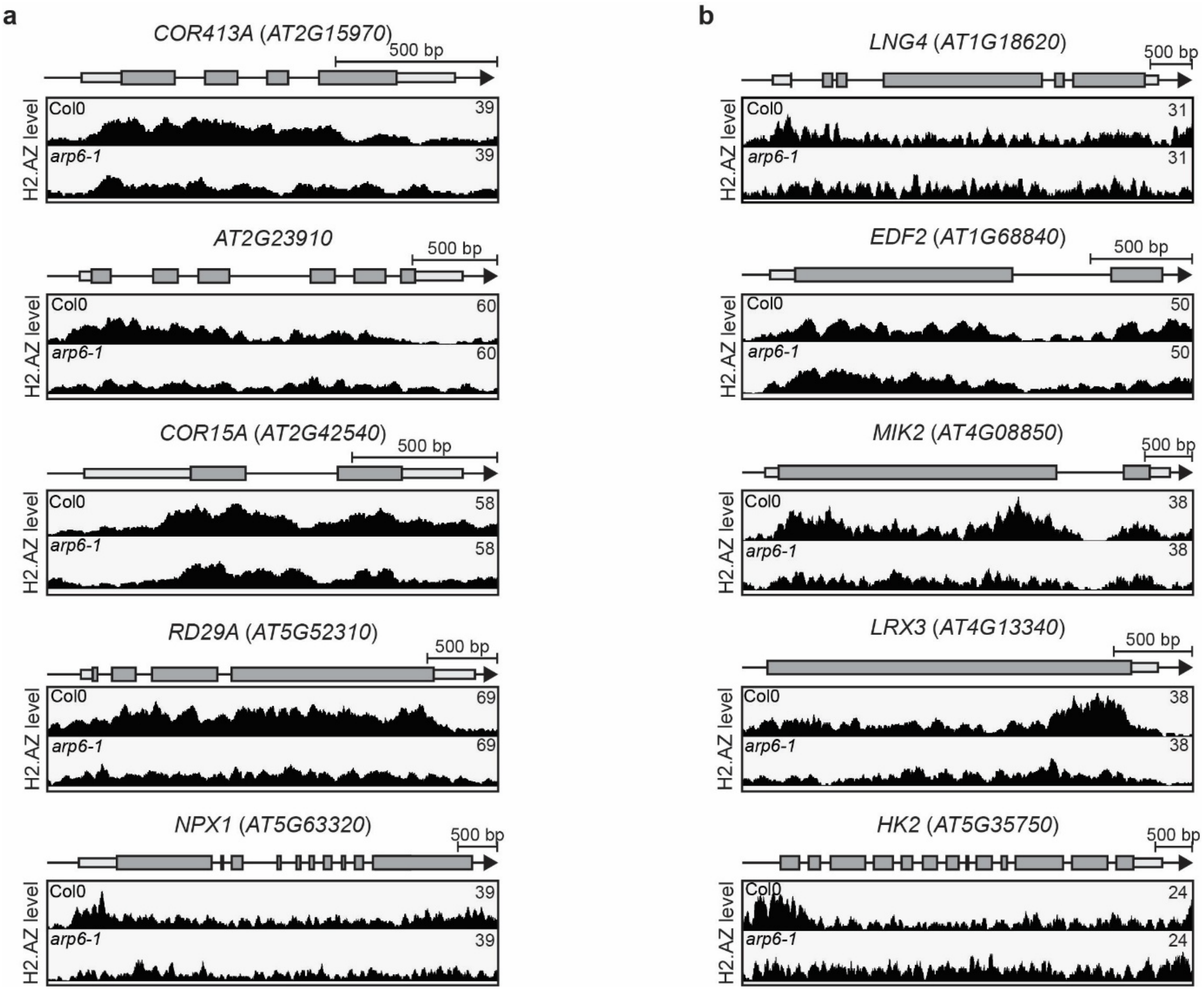


**Supp. Figure 5. H2A.Z profiles in Col0 and *arp6-1* mutant of cold-deregulated genes with cold-enrichment H2A.Z level changes used qPCR analysis**

***a-b)*** H2A.Z profiles at 22°C, from published data[39], in Col0 and *arp6-1* mutant at examples of cold-induced genes with cold-depleted level of H2A.Z (**a**) and cold-repressed genes with cold-enriched level of H2A.Z (**b**) selected for qPCR analysis. Screenshots are from ChIP-seq of H2A.Z in Col0 and *arp6-1* mutant datasets. Higher occupancy of H2A.Z are indicated by higher peak density and amplitude.

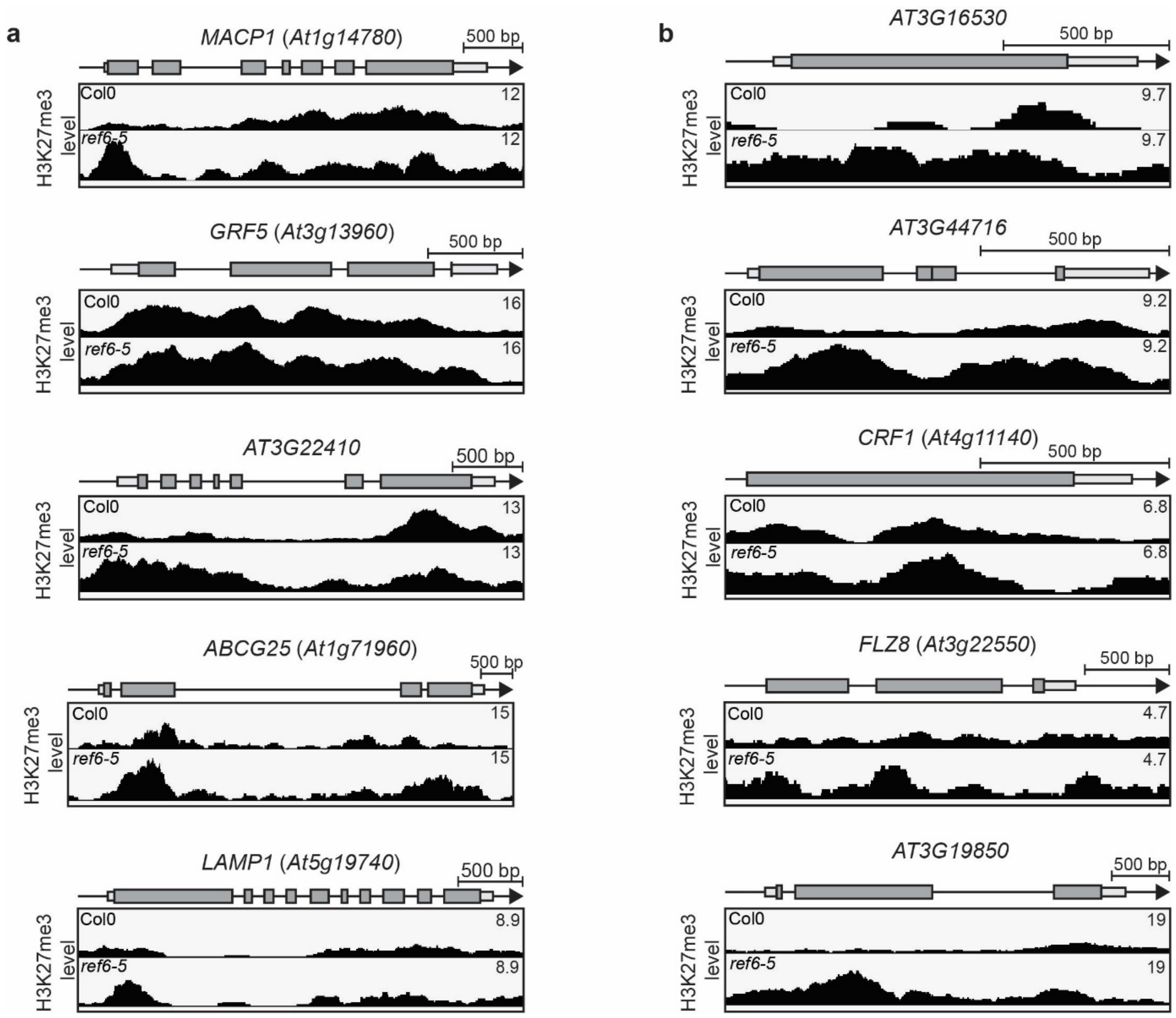


**Supp. Figure 6. H3K27me3 profiles in Col0 and *ref6-5* mutant of cold-deregulated genes with a cold-depleted H3K27me3 level and binding REF6 in their promoter used for qPCR analysis**

***a-b)*** H3K27me3 profiles at 22°C, from published data[43], in Col0 and *ref6-5* mutant at examples of cold-induced genes with cold-depleted level of H3K27me3 (**a**) and cold-repressed genes with cold-depleted level of H3K27me3 (**b**) selected for qPCR analysis. Screenshots are from ChIP-seq of H3K27me3 in Col0 and *ref6-5* mutant datasets. Higher occupancy of H3K27me3 are indicated by higher peak density and amplitude.

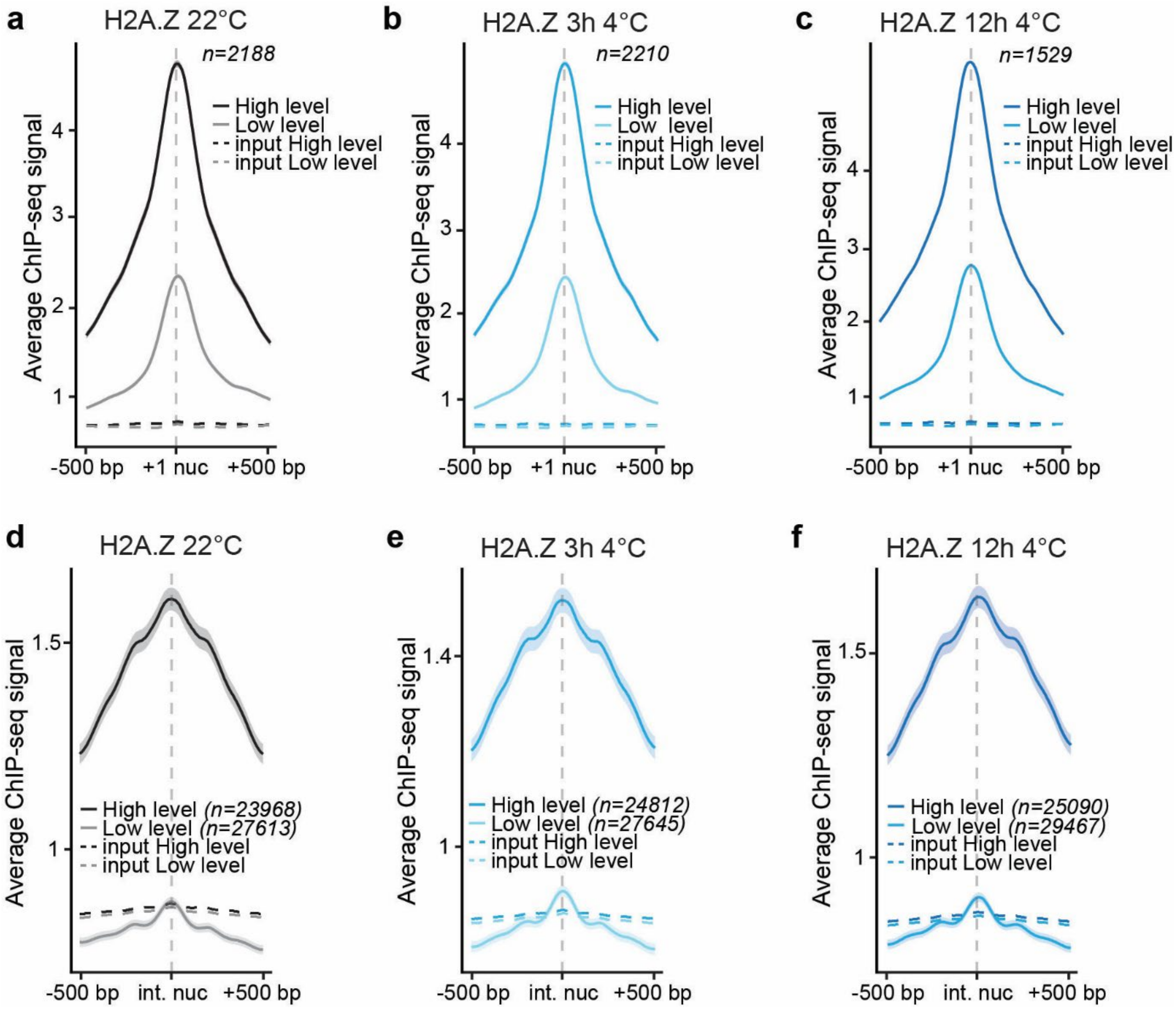


**Supp. Figure 7. Nucleosome groups with low or high levels of H2A.Z.**

**a-f)** Metagene plots of average signal from ChIP-seq of H2A.Z at 22°C (**a, d**), 3h 4°C (**b, e**) or 12h 4°C (**c, f**) of genes classified as high-signal (10% highest, darker lines) or low-signal (10% lowest, lighter lines) in H2A.Z at the +1 nucleosome (first nucleosome downstream the TSS) (**a-c**) or internal nucleosomes (nucleosomes under the peak in the gene body without +1 nucleosome) (**d-f**). Dashed lines indicate the corresponding input signals. The metagene plot is centered on the nucleosome center and includes 500 bp flanks upstream and downstream of the nucleosome center (+1 or internal). The shaded area shows a 95% confidence interval for the mean.

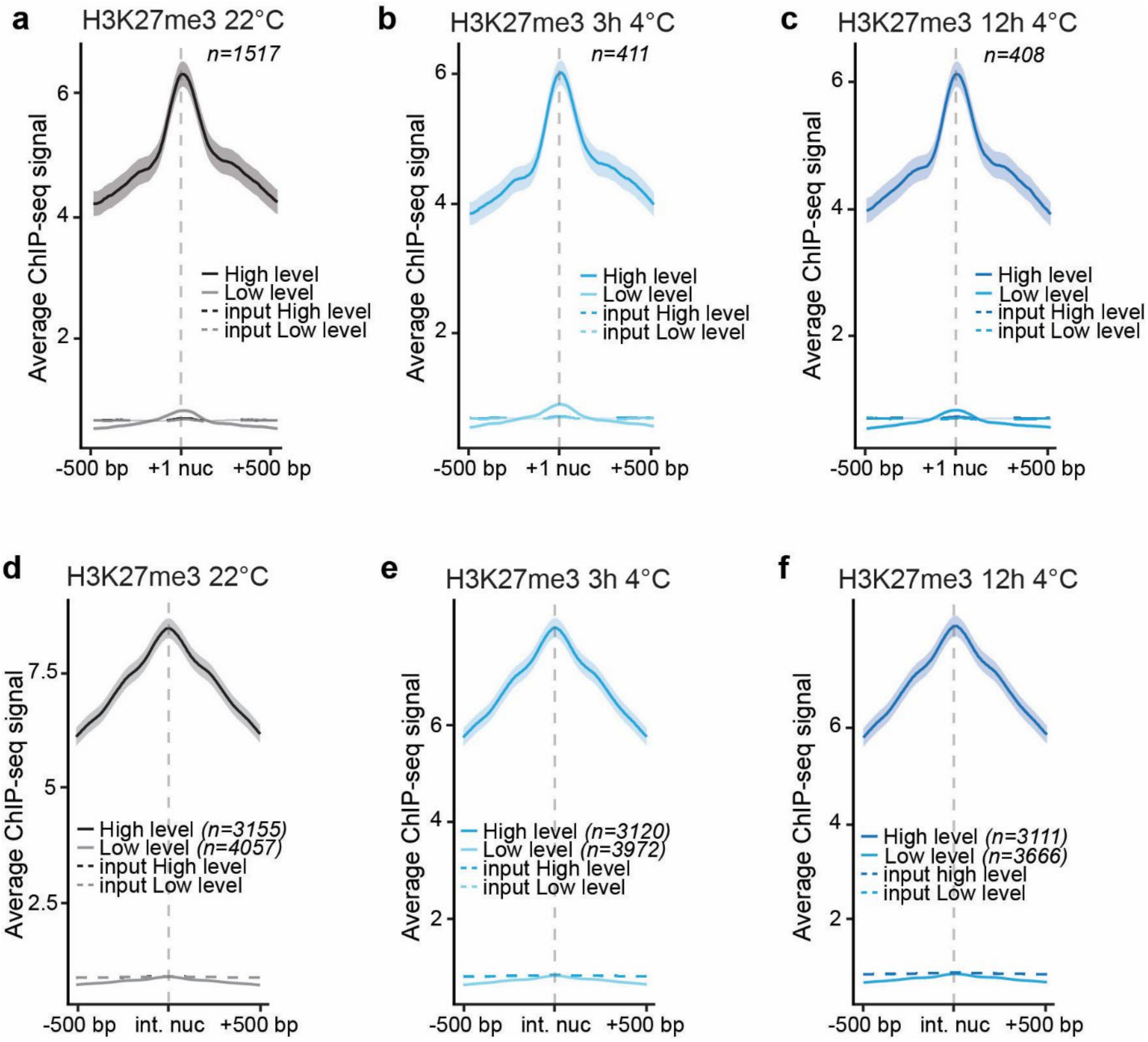


**Supp. Figure 8. Nucleosome groups with low or high levels of H3K27me3.**

**a-f)** Metagene plots of average signal from ChIP-seq of H3K27me3 at 22°C (**a, d**), 3h 4°C (**b, e**) or 12h 4°C (**c, f**) of genes classified as high-signal (10% highest, darker lines) or low-signal (10% lowest, lighter lines) in H3K27me3 at the +1 nucleosome (first nucleosome downstream the TSS) (**a-c**) or internal nucleosomes (nucleosomes under the peak in the gene body without +1 nucleosome) (**d-f**). Dashed lines indicate the corresponding input signals, which largely overlap across conditions and may therefore be difficult to visually distinguish. The metagene plot is centered on the nucleosome center and includes 500 bp flanks upstream and downstream of the nucleosome center (+1 or internal). The shaded area shows a 95% confidence interval for the mean.

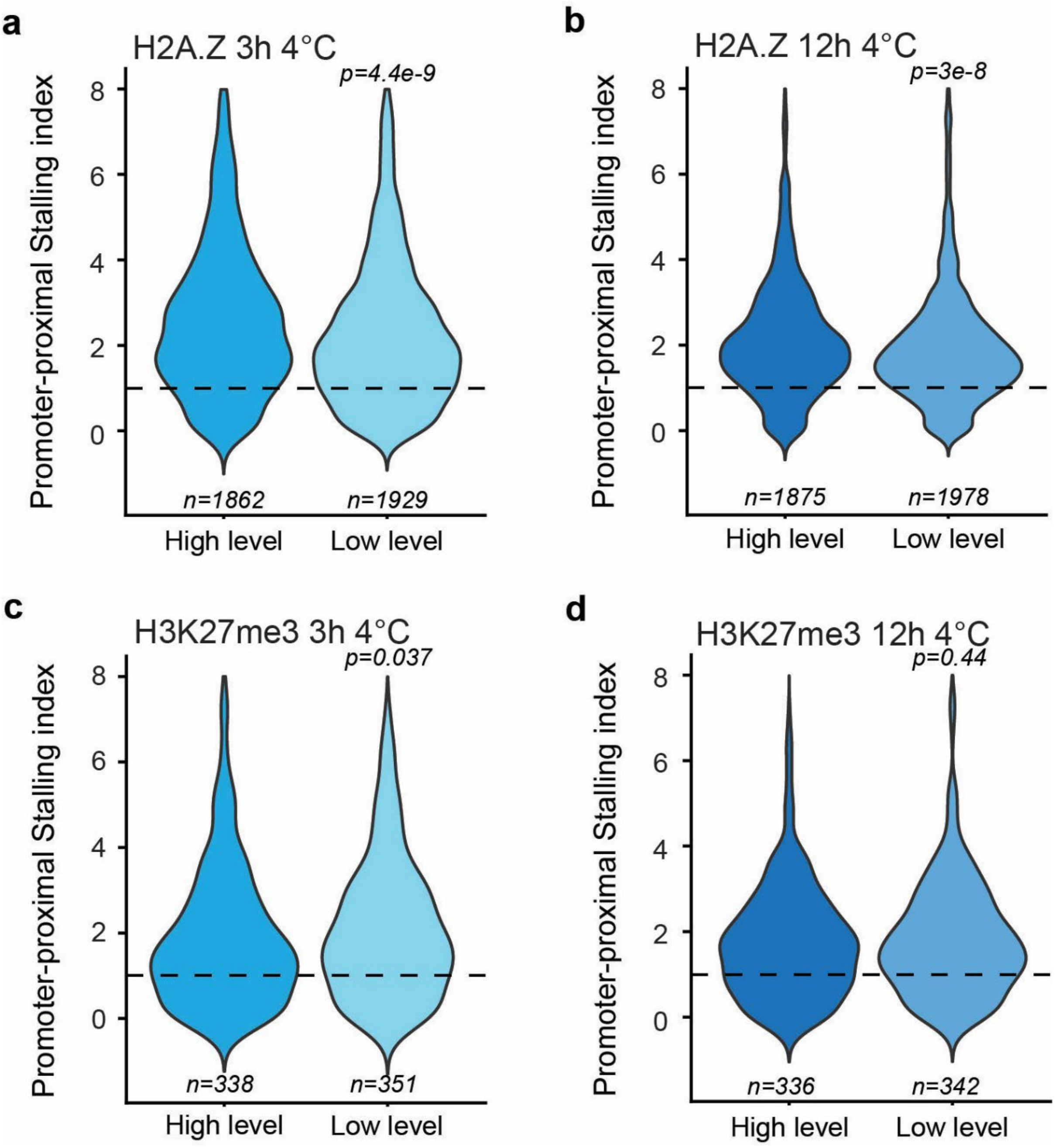


**Supp. Figure 9. Promoter-proximal Stalling Index (PSI) for low and high levels of H2A.Z and H3K27me3.**

**a-d)** Violin plots showing the Promoter-proximal Stalling Index (PSI) of genes with H2A.Z (**a**, **b**) or H3K27me3 (**c, d**) peaks at 3h 4°C (**a**, **c**) or 12h 4°C (**b, d**). Peaks were classified into high-signal and low-signal groups according to the ChIP-seq signal intensity of the corresponding histone mark, corresponding to the top 10% and bottom 10% of peaks, respectively. Dashed lines indicate PSI = 1. Values around 1 indicate no strong RNAPII accumulation, whereas higher values indicate increased RNAPII pausing. p-values indicate statistical significance of differences between conditions, assessed using two-sided unpaired Wilcoxon signed-rank tests, with $p < 0.05$ considered significant.

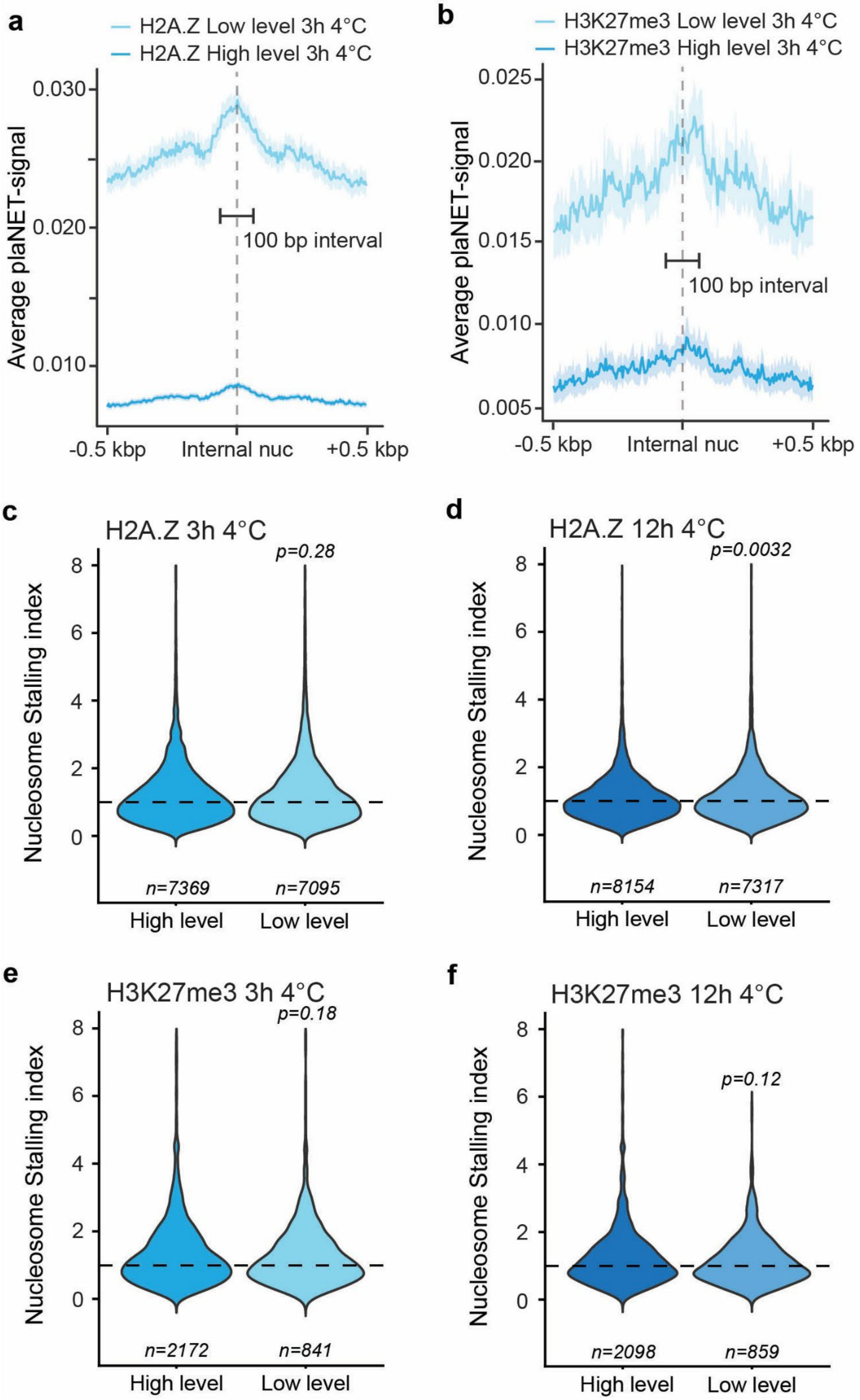
a
H2A.Z Low level 3h 4°C
H2A.Z High level 3h 4°C
Average plaNET-signal
0.030
0.020
0.010
100 bp interval
-0.5 kbp
Internal nuc
+0.5 kbp
b
H3K27me3 Low level 3h 4°C
H3K27me3 High level 3h 4°C
Average plaNET-signal
0.025
0.020
0.015
0.010
0.005
100 bp interval
-0.5 kbp
Internal nuc
+0.5 kbp
c
H2A.Z 3h 4°C
p=0.28
Nucleosome Stalling index
n=7369
n=7095
High level
Low level
d
H2A.Z 12h 4°C
p=0.0032
Nucleosome Stalling index
n=8154
n=7317
High level
Low level
e
H3K27me3 3h 4°C
p=0.18
Nucleosome Stalling index
n=2172
n=841
High level
Low level
f
H3K27me3 12h 4°C
p=0.12
Nucleosome Stalling index
n=2098
n=859
High level
Low level

**Supp. Figure 10. Nucleosome Stalling Index (NSI) for depleted and enriched levels of H2A.Z and H3K27me3.**

**a-b)** Metagene plots of average signal from plaNET-seq of peaks classified as high-signal (10% highest, darker lines) or low-signal (10% lowest, lighter lines) in H2A.Z (**a**) or H3K27me3 (**b**) at the internal nucleosomes (nucleosomes under the peak in the gene body without +1 nucleosome) at 3h 4°C. The metagene plot is centered on the nucleosome center and includes 500 bp flanks upstream and downstream of the internal nucleosome center. The shaded area shows a 95% confidence interval for the mean. The 100 bp interval indicated corresponds to the region used for nucleosome stalling index (NSI) calculation in panels **c** and **e**.

**c-f)** Violin plots showing the nucleosome stalling index (NSI) of genes with H2A.Z (**c**, **d**) or H3K27me3 (**e, f**) peaks at 3h 4°C (**c, e**) or 12h 4°C (**d, f**). Peaks were classified into high-signal and low-signal groups according to the ChIP-seq signal intensity of the corresponding histone mark, corresponding to the top 10% and bottom 10% of peaks, respectively. Dashed lines indicate NSI = 1. Values around 1 indicate no strong RNAPII accumulation, whereas higher values indicate increased RNAPII pausing. p-values indicate statistical significance of differences between conditions, assessed using two-sided unpaired Wilcoxon signed-rank tests, with $p < 0.05$ considered significant.
